# Bottom-up design of calcium channels from defined selectivity filter geometry

**DOI:** 10.1101/2024.12.19.629320

**Authors:** Yulai Liu, Connor Weidle, Ljubica Mihaljević, Joseph L. Watson, Zhe Li, Le Tracy Yu, Sagardip Majumder, Andrew J. Borst, Kenneth D. Carr, Ryan D. Kibler, Tamer M. Gamal El-Din, William A. Catterall, David Baker

## Abstract

Native ion channels play key roles in biological systems, and engineered versions are widely used as chemogenetic tools and in sensing devices^1,2^. Protein design has been harnessed to generate pore-containing transmembrane proteins, but the capability to design ion selectivity based on the interactions between ions and selectivity filter residues, a crucial feature of native ion channels^3^, has been constrained by the lack of methods to place the metal-coordinating residues with atomic-level precision. Here we describe a bottom-up RFdiffusion-based approach to construct Ca^2+^ channels from defined selectivity filter residue geometries, and use this approach to design symmetric oligomeric channels with Ca^2+^ selectivity filters having different coordination numbers and different geometries at the entrance of a wide pore buttressed by multiple transmembrane helices. The designed channel proteins assemble into homogenous pore-containing particles, and for both tetrameric and hexameric ion-coordinating configurations, patch-clamp experiments show that the designed channels have higher conductances for Ca^2+^ than for Na^+^ and other divalent ions (Sr^2+^ and Mg^2+^). Cryo-electron microscopy indicates that the design method has high accuracy: the structure of the hexameric Ca^2+^ channel is nearly identical to the design model. Our bottom-up design approach now enables the testing of hypotheses relating filter geometry to ion selectivity by direct construction, and provides a roadmap for creating selective ion channels for a wide range of applications.

## Main

The origins of the ion selectivity of natural channels has been of great interest since the characterization of the first native channels, but experimental approaches to date have been limited to examining the effects of amino acid substitutions^4–9^ or domain grafting among homologs that creates chimeric channels^10–12^ and have not allowed systematic probing of pore geometry. The ability to construct ion channels from scratch with pre-defined selectivity filters and pore geometries would enable direct testing of hypotheses relating pore structure to conductance, and the creation of custom channels for chemogenetics and sensing applications^1,2,13,14^. Progress has been made in de novo design of pore-containing transmembrane proteins and channels, including assembly of synthetic peptides into barrels^15–19^, conversion of water-soluble helical bundles into transmembrane pores^20,21^, and the design of transmembrane β-barrels from 2D blueprints^22,23^. However, these approaches do not enable precisely placing ion-interacting residues along the ion permeation pathway at angstrom level, and hence designed conducting channels to date have had little ion selectivity. This limitation is particularly significant in the context of designing a Ca^2+^ channel since Ca^2+^ has a nearly identical bare ion radius (∼1 Å) to Na^+^ and a hydrated ion radius (∼4 Å) similar to Mg^2+^ ^24–26^. Native Ca^2+^ channels utilize carboxylate groups to bind Ca^2+^ ions with high affinity likely in a partially dehydrated manner contingent on the geometry of the carboxylate groups^27–32^, which prevents permeation of other ions. Such stringent requirements on the geometry of the selectivity filter are not readily achievable using previous design methods, for which the positions of pore-lining residues are largely determined by the prior placement of the protein backbone.

We sought to develop a general approach for designing ion channels based on ion-coordinating residues that function as the selectivity filter. We chose to focus on Ca^2+^ channels as the mechanism of Ca^2+^ selectivity is relatively well understood, and designed Ca^2+^ channels could serve as useful biological tools due to the critical role of Ca^2+^ as a second messenger. With the hypothesis that the proper spatial arrangement of the carboxyl groups at the pore entrance followed by a wide and well-hydrated pore comprised of residues distant from the axis of the pore would confer both Ca^2+^ permeability and selectivity, we reasoned that Ca^2+^ channels could in principle be constructed by (1) generating selectivity filter configurations by systematically sampling the distance and coordination geometry of carboxylate containing sidechains around a central Ca^2+^ ion, and then generative deep learning methods to (2) build single helices out from the selectivity filter sidechains that define the overall pore shape, and (3) buttress these central pore helices by multiple surrounding helices. This procedure could in principle create channels with custom-defined selectivity filter geometry with high precision, with the overall topology of the protein optimal for supporting the selectivity filter and conducting pore rather than being defined in advance.

### Selectivity filter generation

All known native Ca^2+^ channels utilize negatively charged Glu or Asp residues in the selectivity filter, but in different configurations with different backbone flexibilities. For example, in Ca^2+^ channels within the superfamily P-loop channels where the Glu/Asp residues are situated on a loop, Ca_v_1.1 has the selectivity filter asymmetrically formed by 4 glutamate residues with neighboring oxygen atom distances ranging from 3.3 Å to 5.4 Å^33^, the homotetrameric TRPV6 channel has four Asp residues with diagonal oxygen atom distances of 4.6 Å^34^, while this same diagonal distance in the Ca_v_ab channel (engineered from bacterial Na_v_ab channel) is 8 Å^5^; and the Orai channel (which does not have a P-loop) has six Glu residues on a helix with a diagonal oxygen distance of 8 Å (in the solved open-state structure)^35^.

Considering this ambiguity as well as the commonly observed flexibility of the Glu/Asp sidechains in those native channels, to simplify conformational sampling, we focused on the distance (R_s_) between the Ca^2+^ ion and the C alpha (Cα) atom of the Glu (or Asp) regardless of rotamer which we refer to below as the radius of the selectivity filter (Fig. 1a). We place the Glu/Asp residue in the x-y plane with the Ca^2+^ ion. A second residue with unspecified amino acid identity was then placed at a displacement (h) below the Glu/Asp residue along the z axis at a distance of (R_B_) to the z axis, which was used to define the radius of the pore exit (Fig. 1a). Given a group of sampled values of R_s_ and R_B_, four or six fold symmetry around the z axis to generate the framework of the pore holding the selectivity filter at the pore entrance (Fig. 1b; we focused on C4 and C6 symmetries as these are the commonly observed Ca^2+^ coordination states^25^).

**Fig. 1:**
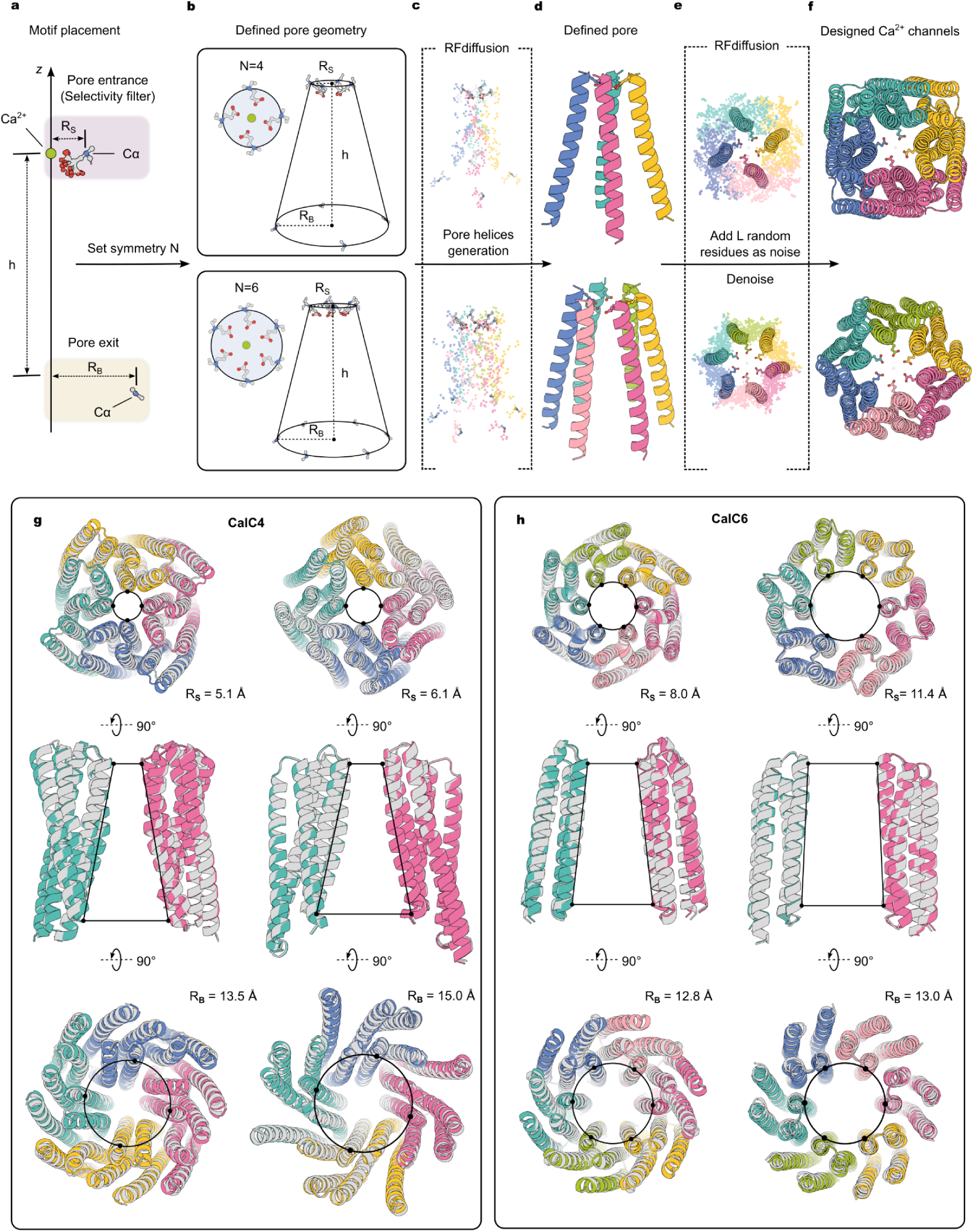
A general approach for designing Ca^2+^ channels from selectivity filter geometry. **a,** A pair of Ca^2+^ ion (green sphere) and an acidic amino acid (Glu in this case, with the C-alpha atom colored in blue) and an additional pore exit defining residue (only backbone atoms shown) are used as the initial structural motifs. The distance between the Ca^2+^ ion and the C-alpha atom (R_s_) defines the size of the selectivity filter regardless of the specific rotamer conformations of the Glu residue. The pore exit defining residue is positioned at a distance (h) along the z-axis from the Glu residue; the distance from the C-alpha atom of this residue to the z-axis (R_B_) defines the radius of the pore exit. **b,** Application of N-fold cyclic symmetry generates the framework of the pore defined by R_s_, R_B_, h and symmetry (N), and the selectivity filter coordinating the Ca^2+^ ion. **c and d,** The selectivity filter and pore exit defining residues are connected by helices using RFdiffusion. **e and f,** The pore-lining helices are extended to form channel proteins using RFdiffusion by adding noise consisting of L residues in each monomer followed by a denoising trajectory. **g and h,** Examples of designed channels (colored models) aligned with AlphaFold2 predicted models (gray) from the top view (top), side view (middle) and bottom view (bottom). For side views only the opposing two chains are shown for clarity. **g**, Two designs with C4 symmetry (CalC4), with backbone RMSD values of 0.8 Å and 0.2 Å to AlphaFold2 predictions, respectively. **h,** Two designs with C6 symmetry (CalC6), with backbone RMSD values of 0.8 Å and 0.5 Å to AlphaFold2 predictions, respectively.

### Supporting scaffold generation and sequence design

Using RFdiffusion^36^, the selectivity filter and exit pore defining residues were connected using helices to form the lining of the homooligomeric pore, and these helices were then extended into multi-secondary structure interacting subunits with L residues in each monomer (Fig. 1c-f). As the topology of the subunits is completely unspecified other than the pore lining helix, the diffusion process generates a wide diversity of solutions. Together, the selectivity filter generation and RFdiffusion scaffolding calculations generate multipass transmembrane protein backbones with central channels and different filter geometries. Sequences were designed using ProteinMPNN^37^ with position-specific amino acid constraints: the selectivity filter residues were fixed at Glu or Asp, the lipid-facing surface residues were constrained to be hydrophobic, and the remaining pore-lining residues were disfavored to be charged to avoid interference with ion permeation through the pore (see Methods). A diverse set of designed channels with varying selectivity filter geometries (the tetramers are denoted CalC4, and the hexamers CalC6) were predicted by AlphaFold2-Multimer^38,39^ to assemble into structures that closely resembled the design models (predicted templating modeling (pTM) score greater than 0.85 and Cα root-mean-square-deviation (RMSD) less than 1 Å, Fig. 1g-h and Extended Data Fig. 1). Designs with many hydrophobic residues extending beyond the defined lipid-embedded regions were discarded, and 23 designs in the CalC4 series, 24 designs in the CalC6 series, and 24 designs in the CalC6_H featuring expanded selectivity filter sizes (H referred to as ‘hollow’) were selected for experimental characterization.

### Experimental characterization of designed Ca^2+^ channels

To identify functional channels, we used a cell-based flux assay with Fura-2 AM as the ion-responsive dye and Ba^2+^ as the surrogate ion for Ca^2+^ to screen for designs showing divalent ion permeability in cells. The genes encoding the designs were inserted into the LentiGuide-BC vector^40^ and were expressed in HEK293T cells by lentiviral transduction to generate a relatively homogeneous expression level of the designed proteins (see Methods). 5 of 23, 6 of 24, and 9 of 24 designs in the CalC4, CalC6 and CalC6_H series, respectively, led to increased Fura-2 fluorescence in response to 2 mM Ba^2+^ compared to the baseline when expressed in cells (Extended Data Fig. 2), consistent with channel formation.

The designs which increased Ba^2+^ flux were expressed in *Escherichia coli* (E. coli), purified by Ni-NTA affinity chromatography from the membrane fraction, and analyzed by size exclusion chromatography (SEC) and negative-stain electron microscopy (ns-EM). 3, 2 and 4 designs in CalC4, CalC6, and CalC6_H series, respectively, eluted as expected for the target oligomeric states in SEC, and formed pore-containing particles on ns-EM grids (Extended Data Fig. 3). CalC4_24, CalC6_3 and CalC6_H4 yielded the most homogeneous particles in each design group when imaged by electron microscopy (Fig. 2). CalC4_24 contains a constriction of ∼1 Å in radius lined by four Glu residues, CalC6_3, a constriction of ∼3 Å lined by six Glu residues, and in CalC6_H4, six Glu are placed at ∼7 Å (Fig. 2a-c). For these designs, the peak elution fraction on SEC (Fig. 2d, ∼ 12.5 mL) yielded homogeneous protein particles of ∼8 nm in diameter evident in ns-EM micrographs and 2D class averages (Fig. 2e), consistent with the design models. Circular dichroism (CD) spectroscopy showed characteristic alpha-helix spectra for all three designs; CD melting experiments suggested these designs did not unfold at 95 °C (Extended Data Fig. 4). AlphaFold3^41^ predictions showed Ca^2+^ ions at the designed sites in CalC4_24 and CalC6_3, albeit with relatively low confidence levels (pLDDT less than 80, Extended Data Fig. 5).

**Fig. 2:**
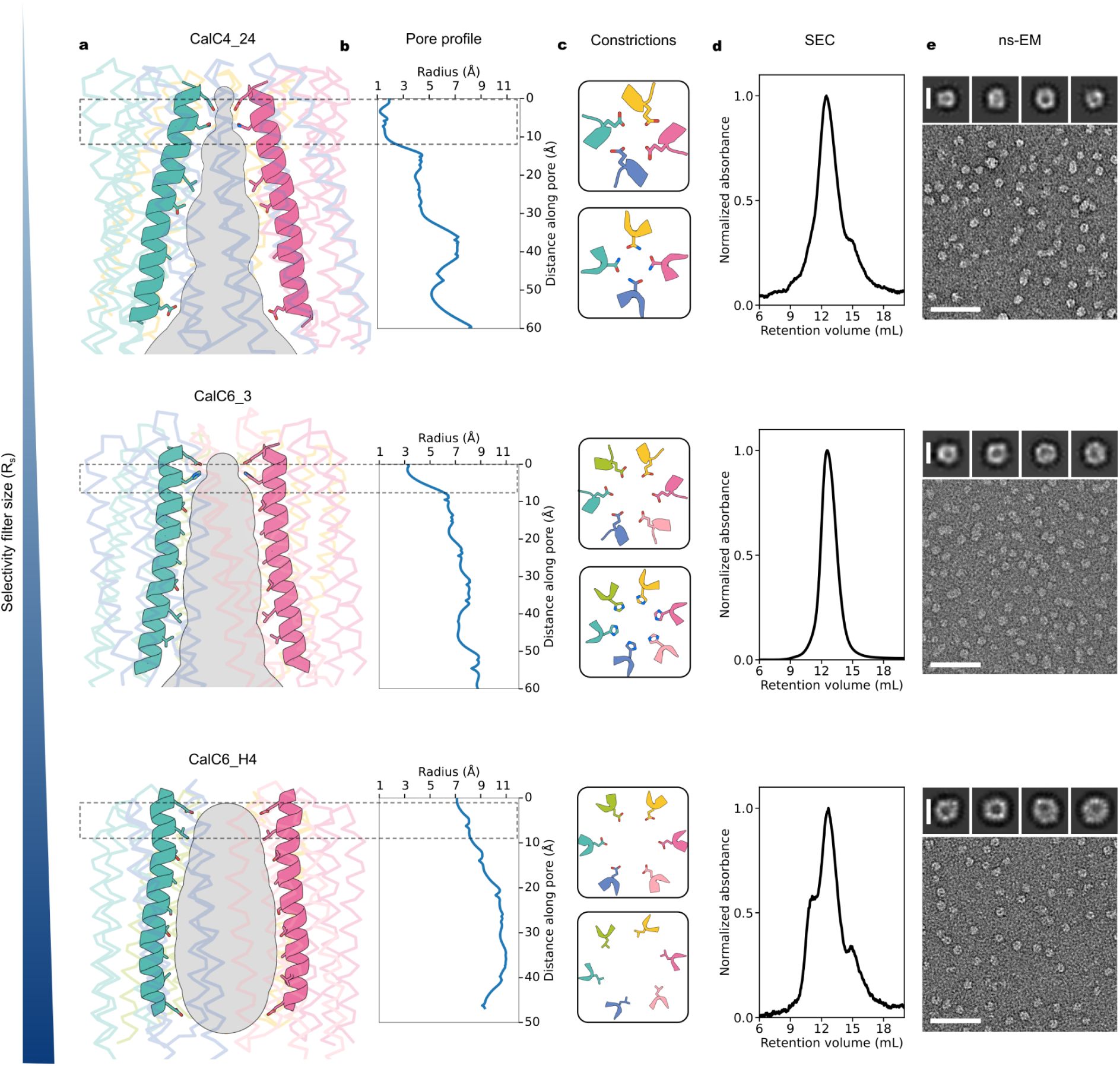
Biophysical characterization of designed Ca^2+^ channels. **a,** Side views of the designed Ca^2+^ channels. Ion permeation pathway (calculated using MOLEonline) and the pore-lining helices on two opposing chains are highlighted, with the remainder of the channels rendered in transparency for clarity. **b,** Pore radius profiles representing the ion permeation pathways in **(a)**. **c,** Top views of the constrictions formed by the sidechains of the residues on two consecutive helical turns. The constrictions are located at the pore entrance, shown by the dashed frame in **(a)** and **(b)**. **d,** Size-exclusion chromatography traces of the purified Ca^2+^ channels. **e,** Representative ns-EM micrographs (scale bars, 50 nm) and 2D class averages (scale bars, 8 nm) of the designed channels.

Next we sought to investigate the conductances of the designed channels for Ca^2+^ by whole-cell patch-clamp recordings on insect *Trichoplusia ni* cells (Hi5) expressing the designs. An experimental challenge is to subtract the background currents (from the leak between the patch pipette and the cell membrane, and from all other channels present in these cells) from the ionic currents for the designed (and constitutively open) channels. We used N-methyl-D-glucamine (NMDG^+^) and methanesulfonate (MeSO_3_^-^) to replace inorganic ions in both extracellular and intracellular solutions (Fig. 3a), bathed the cells in 0.02 mM [Ca^2+^] solution to obtain the baseline currents, and then increased the Ca^2+^ concentration 500 fold to 10 mM [Ca^2+^] solution to preferentially record Ca^2+^ currents (see Methods). With this experimental setting, inward currents at –100 mV were observed to increase for both CalC4_24 and CalC6_3 in 10 mM [Ca^2+^] bath solution. This increase of inward currents was reversible and was diminished when the bath solution was changed back to 0.02 mM [Ca^2+^] solution (Fig. 3b-c). The I-V relations of both channels obtained from both a –100 mV to +100 mV (with a holding potential of 0 mV) voltage step protocol (Fig. 3d-h) and a voltage ramp protocol (Fig. 3i-k) exhibited sharp inward rectifications, as expected based on our experimental design (Ca^2+^ is only present extracellularly and hence can only give rise to inward currents). A current density of approximately 4 pA pF^-1^ at –100 mV in the 10 mM [Ca^2+^] solution was consistently present in cells infected with control baculovirus that did not encode the designs (made from an empty pFastBac-Dual vector), even with 4 mM MgATP included in the patch pipette (Fig. 3l-m and Extend Data Fig. 6). This current was likely the result of a promiscuous endogenous Ca^2+^-permeable channel(s) in Hi5 cells, as it exhibited inward-rectification as well. The current densities in cells expressing CalC4_24 and CalC6_3 were determined to be 7.5 pA pF^-1^ and 9.1 pA pF^-1^, respectively, both significantly larger than the background current (Fig. 3l-m). We did not proceed with testing the CalC6_H4 design by whole-cell patch-clamp experiments due to the possibility of additional ions such as NMDG^+^ or MeSO3^-^ also permeating through the pore, making separation of leak currents more difficult.

**Fig. 3:**
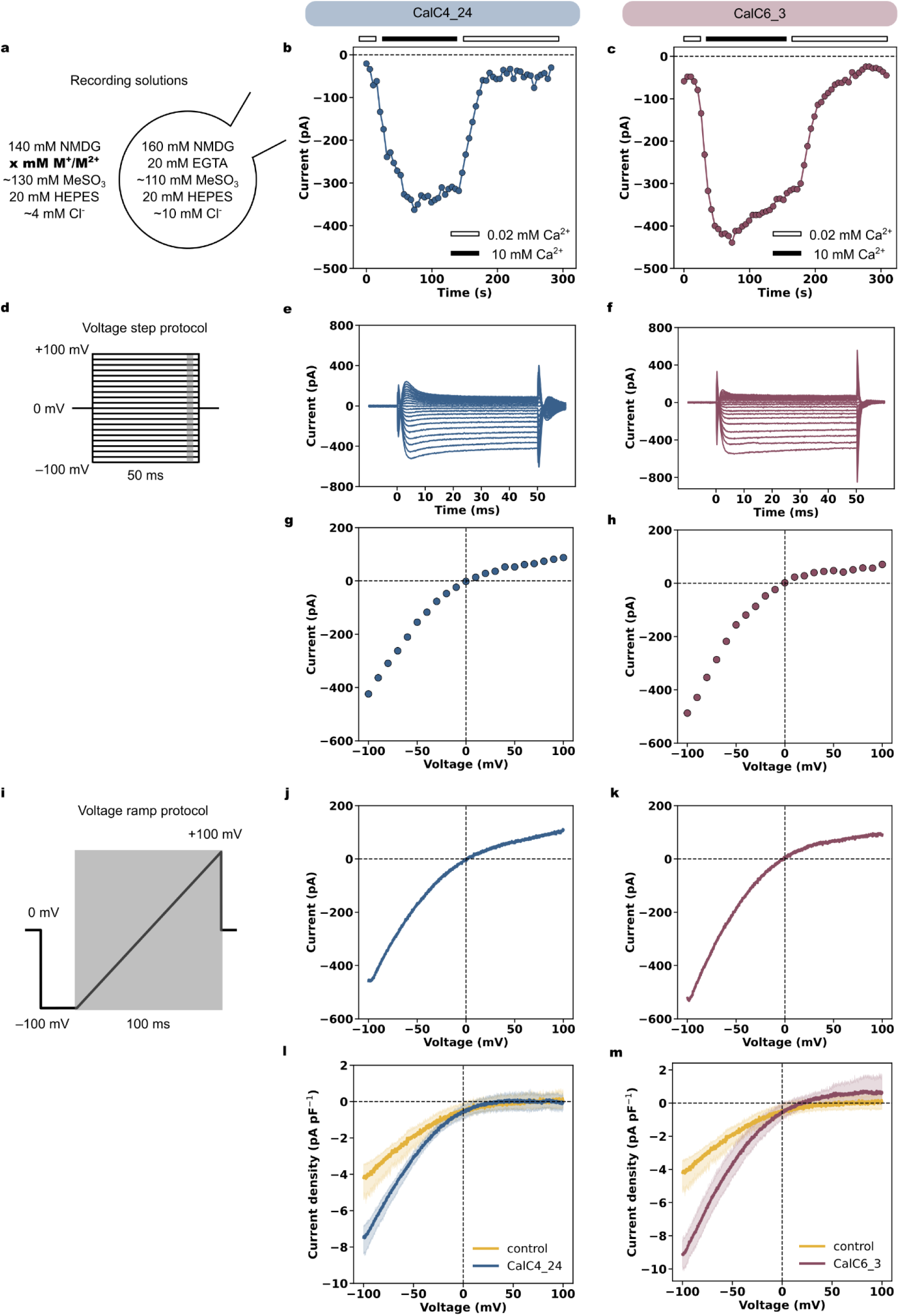
Ca^2+^ conduction by the designed channels. **a,** Diagram showing the solutions used for whole-cell patch-clamp recordings. M^+^ denotes monovalent ions and M^2+^ denotes divalent ions. **b** and **c,** Time courses of the inward whole-cell currents of CalC4_24 **(b)** and CalC6_3 **(c)** recorded at –100 mV in Hi5 cells. Cells were first bathed in 20 µM [Ca^2+^] NMDG-based solution (the first unfilled bars). The extracellular solution was then changed to 10 mM [Ca^2+^] solution (the black bars), and was finally changed back to 20 µM [Ca^2+^] solution (the third unfilled bars). **d,** The schematic illustration of the voltage step protocol. **e** and **f**, Ca^2+^ currents elicited by the voltage step protocol recorded on Hi5 cells expressing CalC4_24 (**e**) and CalC6_3 (**f**), respectively. **g** and **h**, the I-V relations obtained from **e** and **f**, respectively. Tail currents (shown as the gray region in **d**) were used to plot the I-V curves. **i,** The schematic illustration of the voltage ramp protocol. The gray region shows the range used for plotting the I-V curves. **j** and **k**, Ca^2+^ currents elicited by the voltage ramp protocol recorded on Hi5 cells expressing CalC4_24 (**j**) and CalC6_3 (**k**), respectively. **l** and **m**, Current densities from cells expressing CalC4_24 (**l**) and CalC6_3 (**m**) in 10 mM [Ca^2+^] solution after background subtraction using currents recorded in 0.02 mM [Ca^2+^] solution. The control curves (yellow) were obtained from cells infected with the baculovirus that did not encode recombinant proteins (made from an empty pFastBac-Dual plasmid), and were included in both **l** and **m** for comparison. Each I-V curve was obtained by averaging over four measurements on separate cells.

To examine the specificity of the channels for different cations, the 10 mM Ca^2+^ in the extracellular solution was substituted with 40 mM Na^+^ ,10 mM Sr^2+^ or 10 mM Mg^2+^ (the Na^+^ concentration was four times that of the divalent ions as the current scales with the square of the ion valence; see Supplementary information). The relative permeabilities of ions through ion channels are often calculated from the reversal potential using the Goldman-Hodgkin-Katz (GHK) voltage equation when two ions with known concentrations are put on the extracellular and intracellular sides, respectively^42,43^. However, in our case this is complicated by the concentration gradient-driven diffusion through the open channels, which results in variation in the ion concentrations, and the contribution of endogenous channels to the reversal potential is difficult to disentangle. We instead compared conductances for each cation based on the inward current at –100 mV using the GHK flux equation (not the voltage equation, see Methods). Both CalC4_24 and CalC6_3 showed ∼5 fold larger current in 10 mM [Ca^2+^] than in 40 mM [Na^+^] solution at –100 mV (Fig. 4a-d, Extended Data Fig. 6). The relative conductances of the divalent cations for both designs followed the order Ca^2+^ > Sr^2+^ > Mg^2+^ (Fig. 4e-h), as observed for several native Ca^2+^ channels (Ca_v_, TRPV6, CRAC)^30,44–49^. Likewise, both channels were blocked by 10 µM La^3+^, which blocks native Ca^2+^ channels (Fig 4i-l).

**Fig 4:**
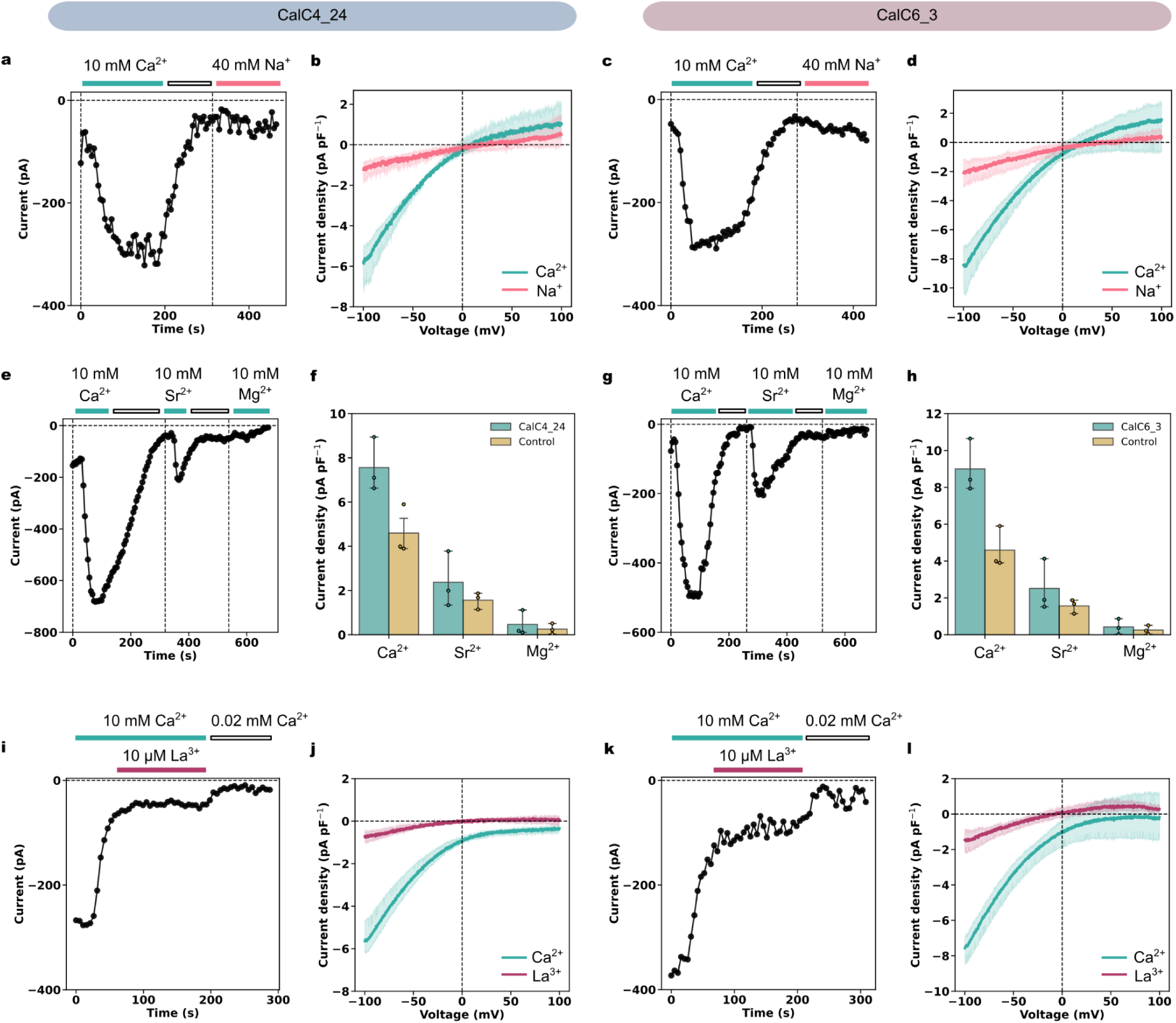
Ion selectivity of designed Ca^2+^ channels. **a-d,** Relative conductances of CalC4_24 **(a-b)** and CalC6_3 **(c-d)** for Ca^2+^ and Na^+^. **a** and **c**, Time courses of currents at –100 mV. Extracellular solution was exchanged from a solution containing 10 mM [Ca^2+^] to 40 mM [Na^+^] with 0.02 mM [Ca^2+^] solution between the exchange. **b** and **d**, I-V relations obtained from a –100 mV to +100 mV ramp protocol (Ca^2+^ currents in cyan and Na^+^ currents in red). **e-h,** Relative conductances of CalC4_24 **(e-f)** and CalC6_3 **(g-h)** for Ca^2+^, Sr^2+^ and Mg^2+^. **e** and **g**, Time courses of currents at –100 mV. Extracellular solution was exchanged from a solution containing 10 mM [Ca^2+^] to 10 mM [Sr^2+^] and 10 mM [Mg^2+^], with 0.02 mM [Ca^2+^] solution between each exchange. **f** and **h**, Averaged peak current densities at –100 mV in each 10 mM divalent ion solution (n=3). The data obtained from cells infected by the control virus were presented in yellow (n=3) and were included in both plots for comparison. **i-l,** Blockade of Ca^2+^ current of CalC4_24 **(i-j)** and CalC6_3 **(k-l)** by 10 µM La^3+^. **i** and **k**, Time courses of currents at –100 mV recorded in 10 mM [Ca^2+^] solution, followed by addition of 10 µM La^3+^, and finally in 0.02 mM [Ca^2+^] solution. **j** and **l**, I-V relations obtained from a –100 mV to +100 mV ramp protocol before (cyan) and after (purple) addition of 10 µM La^3+^ to the 10 mM [Ca^2+^] solution. Each I-V curve was averaged over three measurements on separate cells, with the baseline current measured in 0.02 mM [Ca^2+^] subtracted.

Both CalC4_24 and CalC6_3 exhibit electrophysiological properties expected for Ca^2+^ channels: inward rectification in presence of extracellular Ca^2+^, higher relative conductances to Ca^2+^ than to other cations, and block by lanthanum. Thus, our approach of arranging Glu/Asp carboxylate residues around a central pore generates Ca^2+^ dependent channels as intended. Although the selectivity filter of CalC6_3 (6.4 Å) is much wider than that in CalC4_24 (2.5 Å, both distances represent that between diagonally opposed oxygen atoms), the electric field strength near the filter is probably stronger in CalC6_3 due to the presence of two extra Glu residues, which likely facilitates recruitment of Ca^2+^ ions. The adjacent Glu residues are predicted by AlphaFold3 to interact and partially dehydrate the Ca^2+^ ion (Extended Data Fig. 5a-b), as in the selectivity filter configuration observed in the crystal structure of the Ca_v_ab channel in complex with dihydropyridine compounds^32^. In our experiments 4 mM MgATP (free [Mg^2+^] estimated to be approximately 400 µM in the presence of 20 mM EGTA^50^) was included in the patch pipette to inhibit a large endogenous non-selective cation current, and this could also reduce the currents observed for Na^+^ and other divalent ions (AlphaFold3 also predicts some binding of Mg^2+^ to the selectivity filter; Extended Data Fig. 5c-d), consistent with observations that Mg^2+^ can bind to selectivity filters but not readily permeate in some native Ca^2+^ channels^6,49,51^. The relative conductances order Ca^2+^ > Sr^2+^ > Na^+^ of the designs likely reflects a preferential binding of ions at the selectivity filter with a high field strength (HFS). To verify the importance of the negative charge at this site, we mutated the Glu residues at the selectivity filter of CalC6_3 design to hydrophobic Leu residues and measured the Ca^2+^ conductance. Mutation of the Glu residues reduced the Ca^2+^ conductance of CalC6_3 channel to the same level as that of the negative control (∼ 4 pA pF^-1^ at –100 mV, Extended Data Fig. 6). The size of the orifice formed by the Leu residues was predicted to be almost identical to that formed by the Glu residues, suggesting that the reduction of Ca^2+^ conductance was most likely a result of the removal of the negative charge at the pore entrance.

### Cryo-EM structure determination

To assess the design accuracy, we sought to determine the structure of CalC6_3 using cryo-electron microscopy (cryo-EM). CalC6_3 in detergent particles appeared homogeneous after being frozen on the grid (Extended Data Fig. 7b) and 4,644 movies were collected; however, top views of the particles could not readily be separated from side views during 2D classification due to the spherical shape of the detergent-bound membrane protein, a well-known challenge for structural determination of relatively small membrane proteins without obvious soluble domains^52^. To facilitate structural determination, we applied a fusion protein strategy: we connected structurally-validated designed helical repeat (DHR) proteins^53^ to the C-terminus of CalC6_3 via straight helices, with linker helices generated using RFdiffusion and sequences designed using ProteinMPNN (see Methods, Extended Data Fig. 7e). Among the six tested channel-DHR fusion proteins, one showed quite homogeneous particles on the ns-EM grid.

We proceeded with cryo-EM data collection with this construct, and obtained a three-dimensional reconstruction to a global resolution of 3.75 Å with C6 symmetry (Fig. 5a and e), and were able to build a structure model based on the density map (Fig. 5b and f). The density for the DHR extension domains were not well resolved possibly due to structural flexibility. We were also able to identify some minor heptamer and octamer species from the 2D classification, and obtained 3D reconstructions for the heptamer to a overall resolution of 4.62 Å with C7 symmetry. (Extended Data Fig. 8). The existence of multimeric proteins in an additional oligomeric state has been observed in naturally-existing proteins, for example calcium homeostasis modulator (CALHM) family members^54,55^.

**Fig 5:**
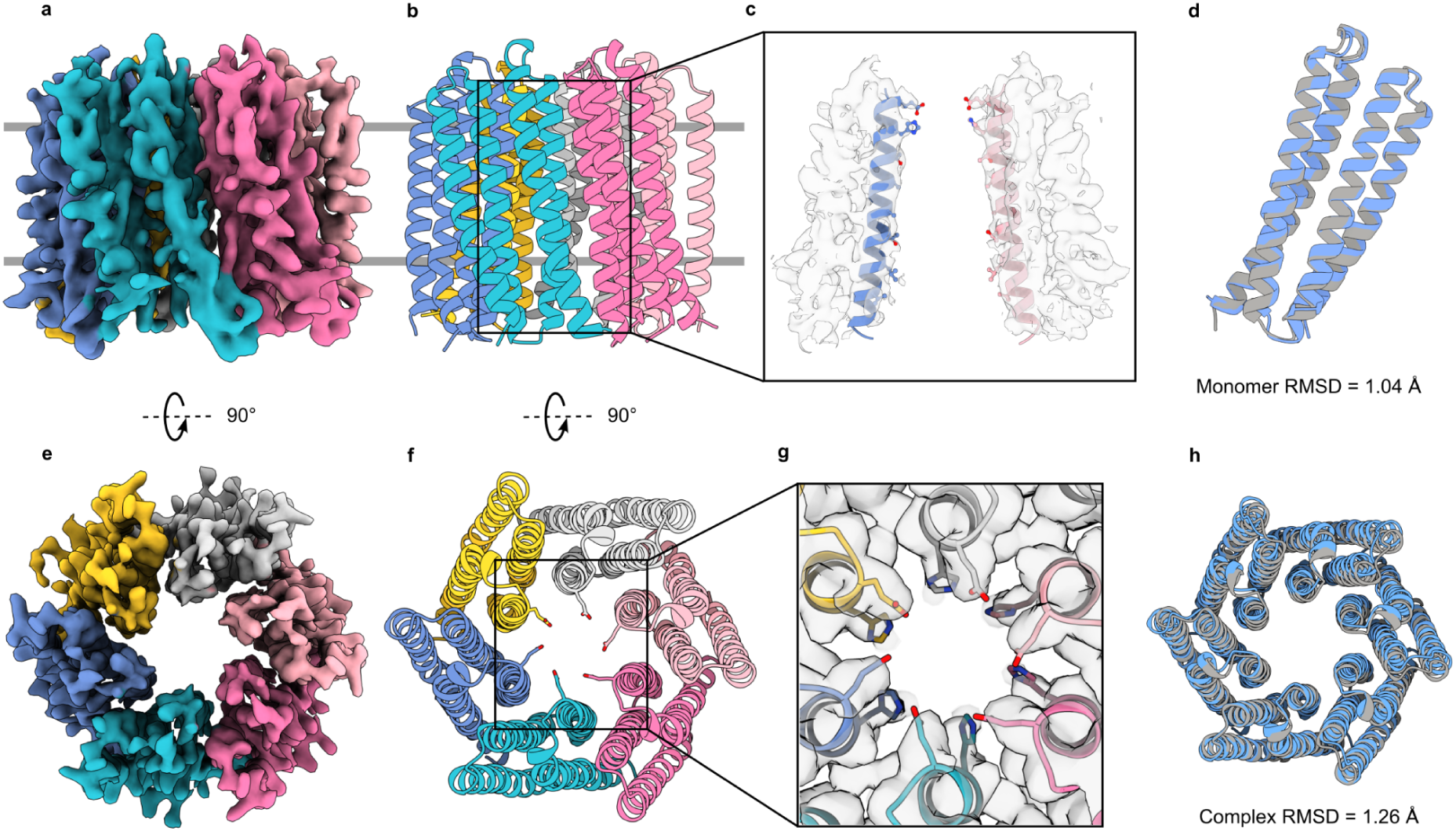
Cryo-EM structure of CalC6_3. **a and e,** Side **(a)** and top **(e)** views of sharpened cryo-EM density map of CalC6_3. **b and f**, Structural model of CalC6_3, with the same colors and orientations as the density map in **(a)** and **(e)**, respectively. **c and g,** Close-up views of the pore-lining helices **(c)** and the selectivity filter **(g)**, with the densities shown as gray surfaces contoured at 4 σ. For the side view of pore helices in **(c)** only the opposing monomers were shown for clarity. **d and h,** Structural alignment of the monomer **(d)** and the hexamer **(h)** of the CalC6_3 design model (gray) to the cryo-EM structure (blue), with backbone RMSDs of 1.04 Å and 1.26 Å for the monomer and complex alignment, respectively.

In the cryo-EM structure of the hexamer, the density for the Glu-ring selectivity filter density and the His-ring below the filter was clearly evident when contoured at and below 4 σ level (Fig. 5c and g), suggesting pre-organization of Glu-ring by the His-ring via potential hydrogen bond interactions. Other than a slight swelling of the cryo-EM complex structure compared to the design model (backbone RMSDs of 1.04 Å and 1.26 Å for monomer and complex alignment, respectively; Fig. 5d and h), the geometry of the selectivity filter was almost exactly as designed.

## Discussion

De novo design of ion channels is in principle a powerful approach to study ion channel biophysics and to develop new biological tools, but the ability to tune ion selectivity has been limited by lack of an approach to install selectivity filters with specified composition and geometry into designed channels. We demonstrate that using generative RFdiffusion symmetric motif scaffolding, protein topologies can be built from scratch to optimally scaffold selectivity filters with pre-defined geometries. In contrast, previously design efforts sought to change conductances by altering the number of alpha helices or beta strands surrounding the pore, rather than building up from defined selectivity filter geometries and compositions. Ca^2+^ selective channels were not obtained using these previous approaches, which is expected given the challenge of selectively recognize Ca^2+^ ions while maintaining a rapid Ca^2+^ ion flow. In contrast, the designed CalC4_24 and CalC6_3 channels have precisely designed selectivity filter geometries, and have higher conductances for Ca^2+^ than for other cations. The high degree of agreement between the cryo-EM structure and the design model of CalC6_3 demonstrates the accuracy of the design method. Our approach enables the exploration of diverse selectivity filter geometries and chemical compositions for optimal ion channel conductance and selectivity, which are difficult to probe through classical native ion channel mutagenesis experiments: for example, it will be very interesting to explore how increasing selectivity filter complexity, and incorporating additional filter layers, impacts channel selectivity. Building up channels around defined selectivity filter geometries should both enable rigorous testing of our understanding of the determinants of native ion channel selectivity, and the construction of a wide array of new channels with selectivities that go beyond those observed in nature.

## Methods

Computational and experimental methods are provided in the Supplementary information.

### Data availability

EM maps of CalC6_3 with DHR extensions have been deposited in the Electron Microscopy Data Bank. Cryo-EM structure files of CalC6_3 with DHR extensions have been deposited to the Protein Data Bank.

### Code availability

Documentation for RFdiffusion is available at https://github.com/RosettaCommons/RFdiffusion. Documentation for ProteinMPNN and LigandMPNN are available at https://github.com/dauparas/ProteinMPNN and https://github.com/dauparas/LigandMPNN, respectively. AlphaFold2 and AlphaFold3 used for filtering the designs are available at https://github.com/google-deepmind/alphafold and https://golgi.sandbox.google.com/ respectively. Example design scripts will be made available on GitHub.

## Acknowledgements

We thank G. R. Lee, N. Hanikel, D. Juergens, G. Wisedchaisri, M. Lenaeus, L. Wang for helpful discussions; D. Feldman for support in lentivirus preparation and transduction; D. E. Clapham and R. E. Hulse for invaluable advice on channel biophysical characterization and comments on the manuscript; E. Navaluna, M. Ahlrichs, S. Cheng, C. Dobbins, for support in tissue culture; J. Quispe and S. Dickinson for management of the Arnold and Mabel Beckman Cryo-EM Center at the University of Washington and the cryo-EM facility at HHMI Janelia Research Campus for cryo-EM usage; and K. VanWormer, L. Goldschmidt and J. Li, for general technical support. This research was supported by the Audacious Project at the Institute for Protein Design (Y.L., R.D.K., and D.B.), National Institutes of Health research grant R01 HL112808 (W.A.C.), the Air Force Office of Scientific Research (FA9550-21-S-0001, Y.L., L.M., S.M. and D.B.), the HHMI Helen Hay Whitney Fellowship (L.M.), and the Howard Hughes Medical Institute (L.M and D.B.). We would also like to express our sadness on the passing of Dr. W. A. Catterall who provided invaluable guidance and inspiration for this work.

## Extended Data

**Extended Data Fig. 1:**
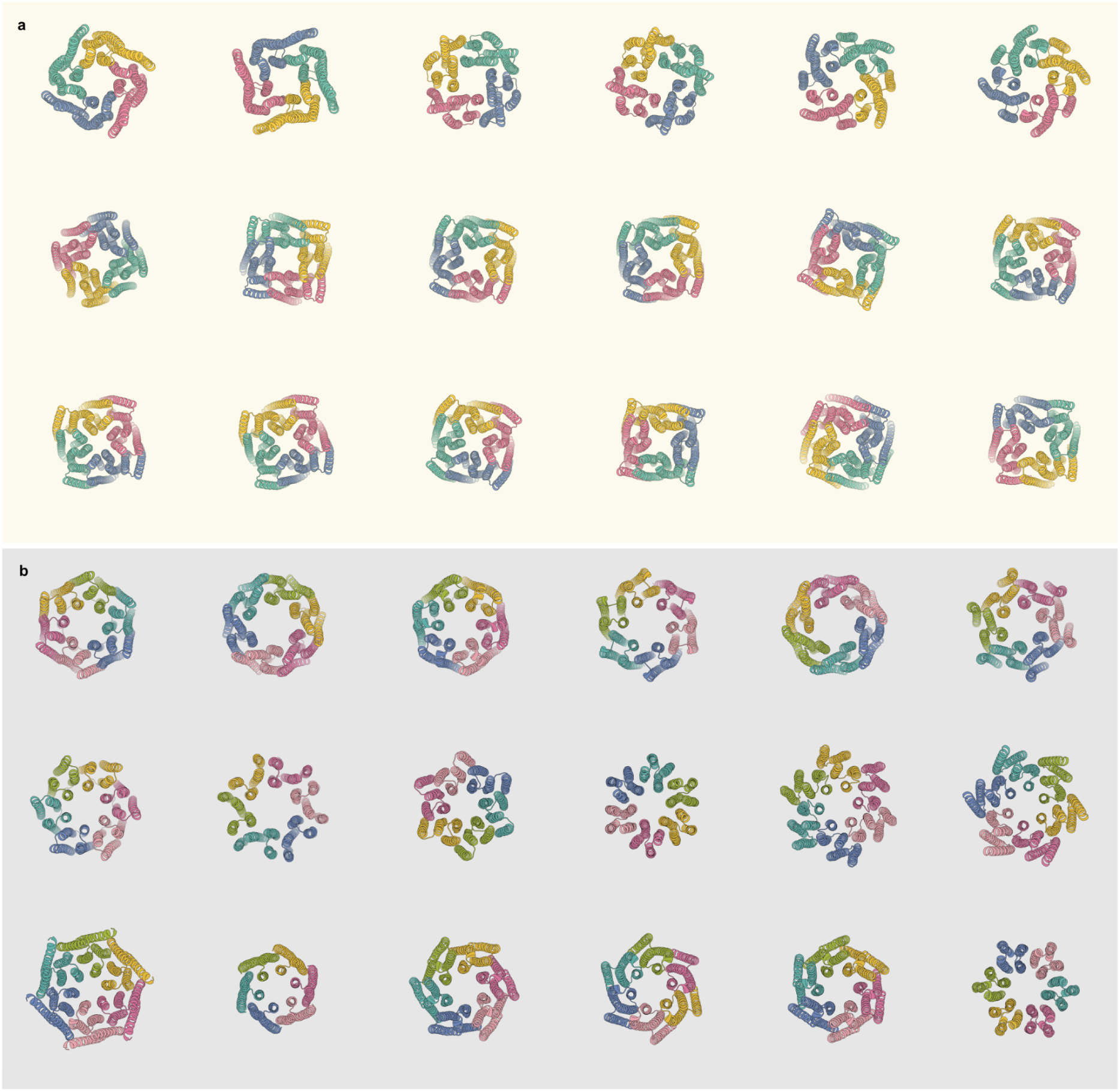
Examples of AlphaFold2-predicted structures of designed Ca^2+^channels with pTM scores higher than 0.85. **a,** Designs with C4 symmetry. **b,** Designs with C6 symmetry.

**Extended Data Fig. 2:**
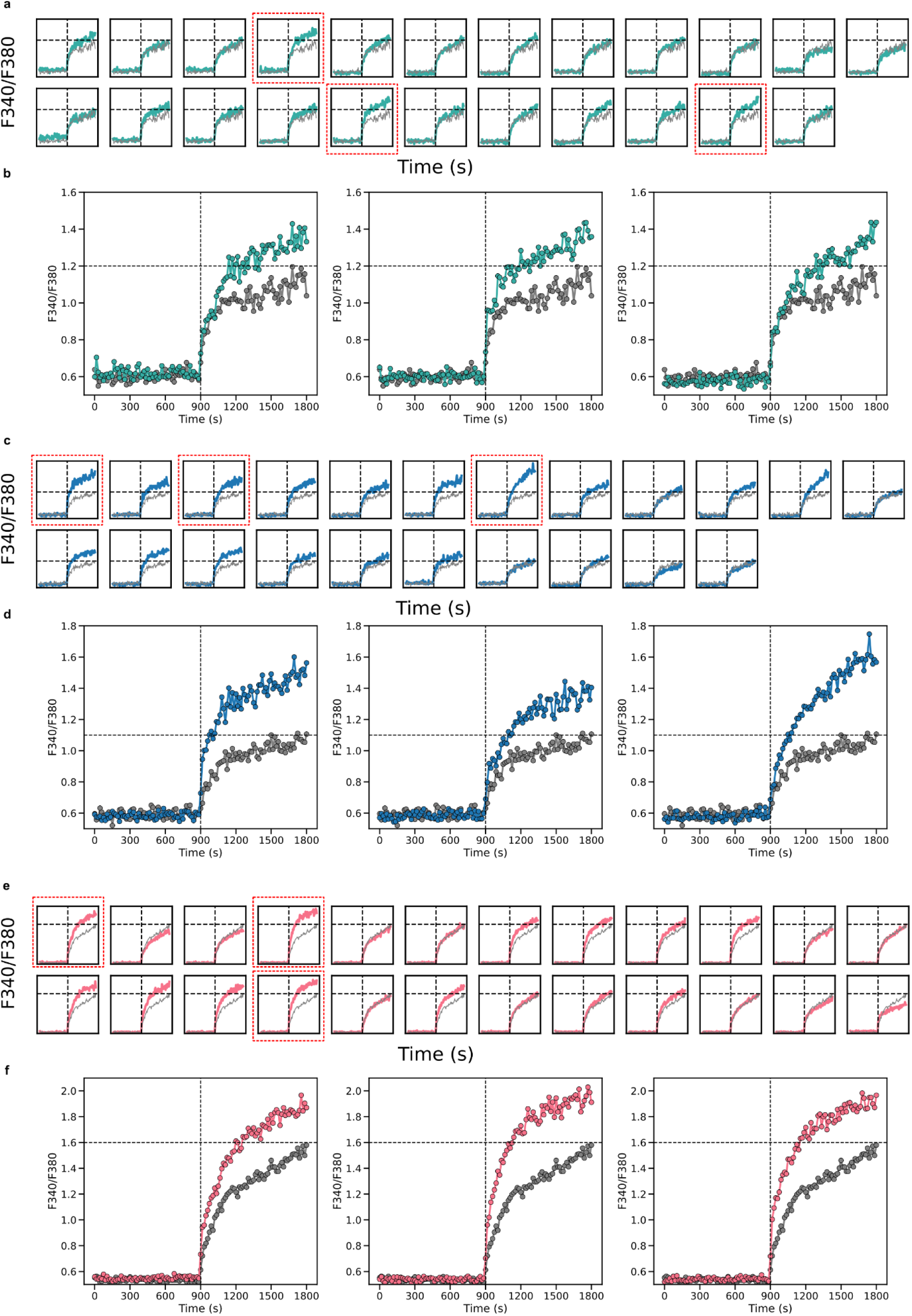
Flux assay in HEK293T cells using Fura-2 AM. **a-b** (cyan), **c-d** (blue), and **e-f** (red) were data for CalC4, CalC6, CalC6_H designs, respectively. Ba^2+^ ions were added at the 900-second time point indicated by the vertical dashed lines. The gray traces represented measurements from mock-transduced cells as the negative control and were overlaid with the colored traces representing the cells transduced with the designs in each plot for comparison. **b**, **d** and **f** were close-up views of the plots highlighted by the red dashed frames in **a**, **c** and **e,** respectively, which were examples that were identified as the putative designs manifesting divalent ion permeability.

**Extended Data Fig. 3:**
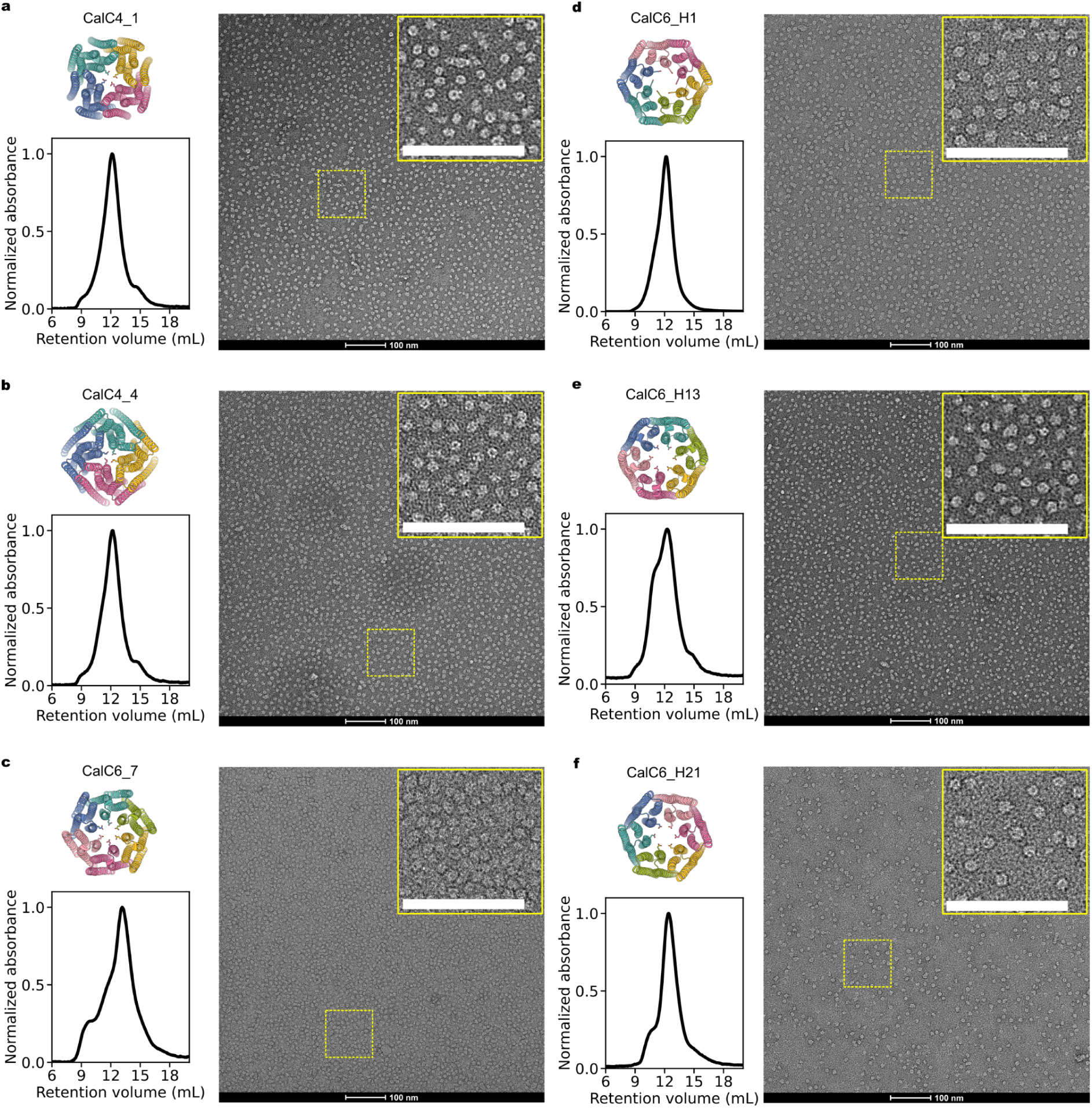
Biophysical characterizations of more designed channels. In addition to the three designs shown in Fig. 2, two more CalC4 designs, CalC4_1 **(a)** and CalC4_4 **(b)**, one more CalC6 design, CalC6_7 **(c)**, and three more CalC6_H designs, CalC6_H1 **(d)**, CalC6_H13 **(e)**, and CalC6_H21 **(f)**, were observed to elute as desired oligomeric states from SEC and assemble into homogeneous pore-containing particles on the ns-EM grids. The close-up views shown on the upper right corners of the ns-EM micrographs corresponded to the regions highlighted by the dashed yellow frames. Scale bars, 100 nm.

**Extended Data Fig. 4:**
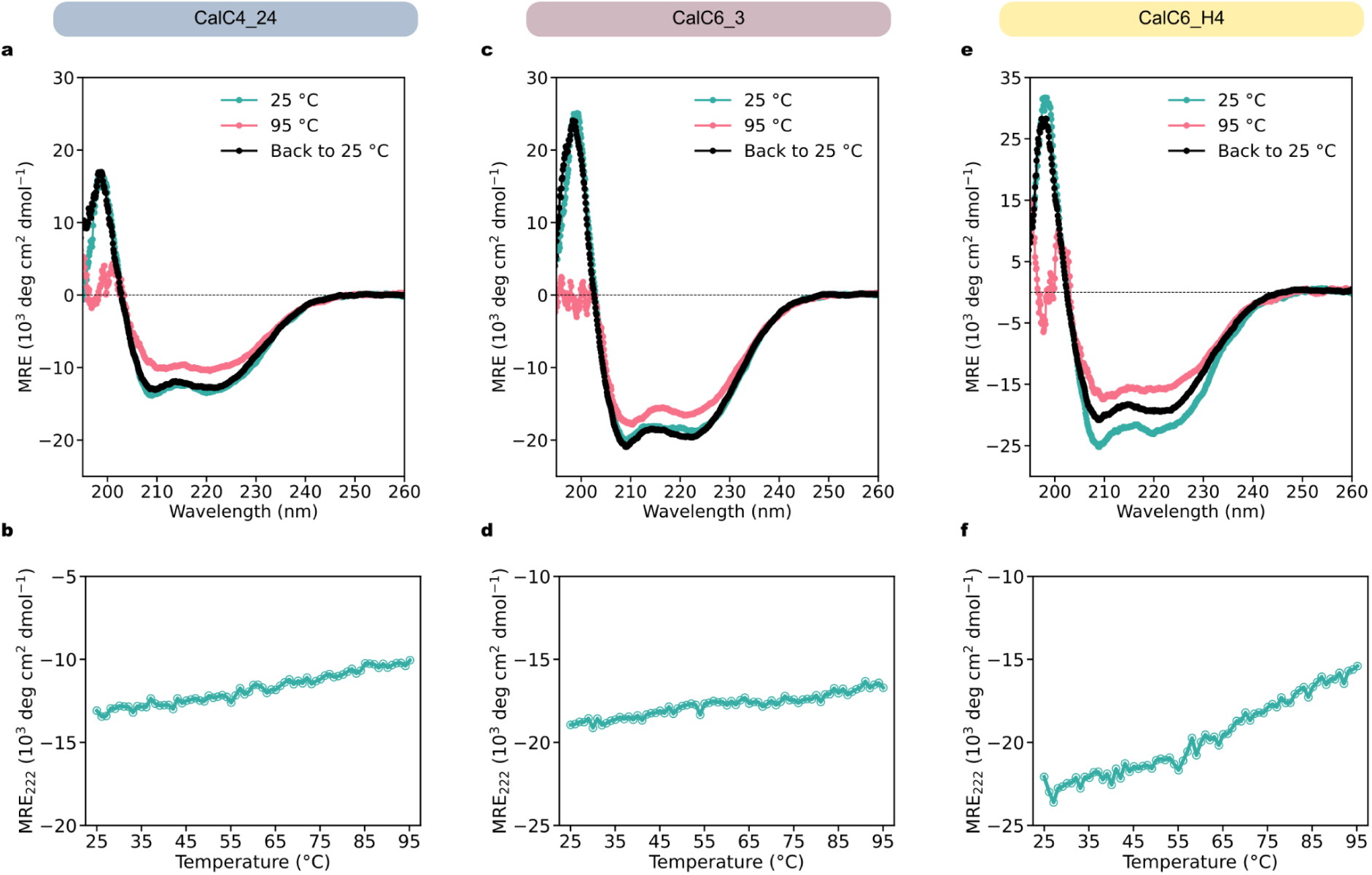
Circular dichroism results of CalC4_24 (**a-b**), CalC6_3 (**c-d**), and CalC6_H4 (**e-f**). **a**, **c**, and **e**, CD spectra at 25 °C (cyan lines), 95 °C (red lines), and cooled back to 25 °C (black lines). **b**, **d**, and **f**, CD melting curves at 222 nm.

**Extended Data Fig. 5:**
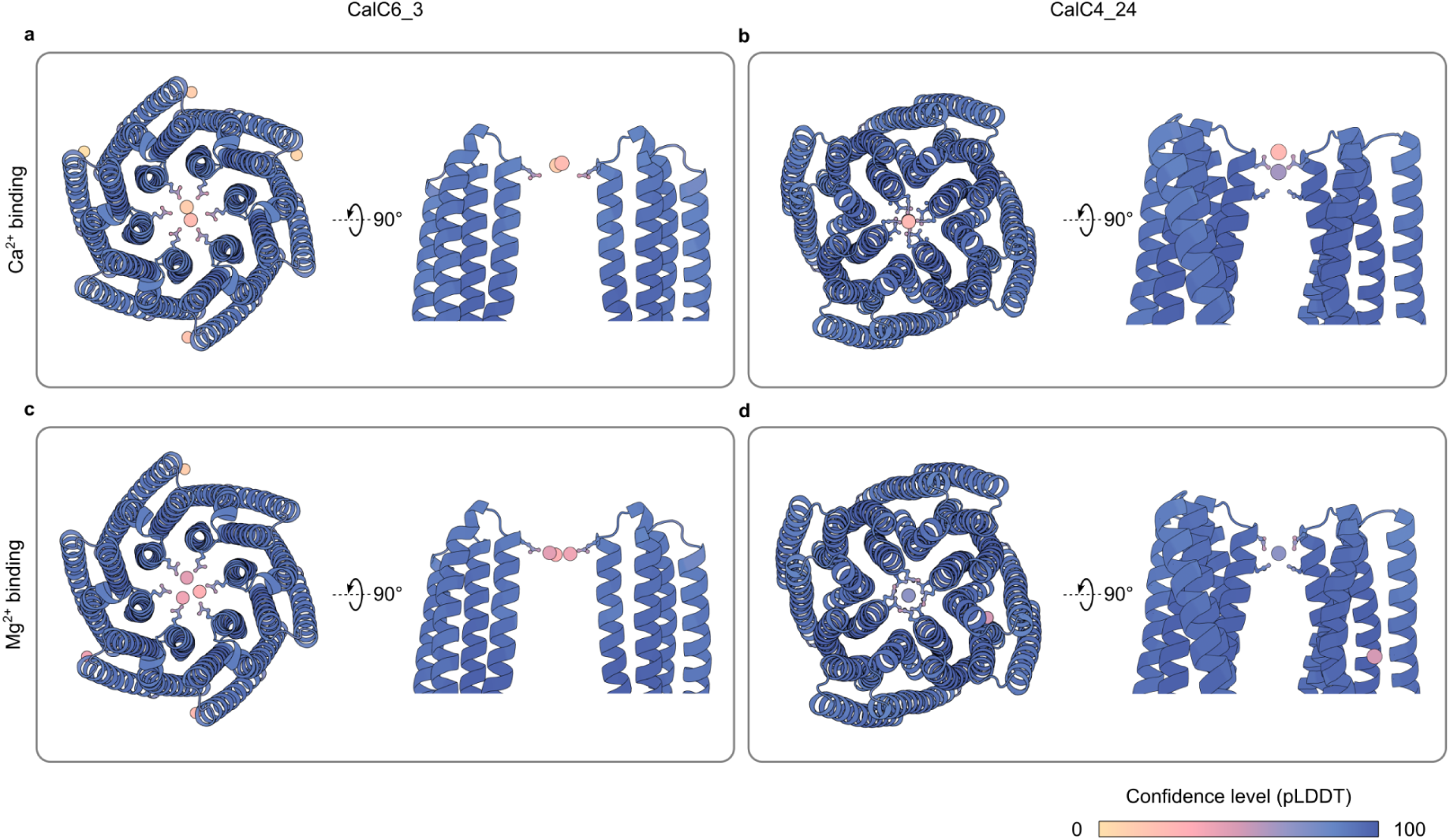
AlphaFold3-predicted binding of CalC6_3 and CalC4_24 binding to Ca^2+^ (**a** and **b**) and to Mg^2+^ (**c** and **d**), respectively. The models were colored based on the confidence level (pLDDT) of the prediction results, with the color gradient shown on the bottom right.

**Extended Data Fig. 6:**
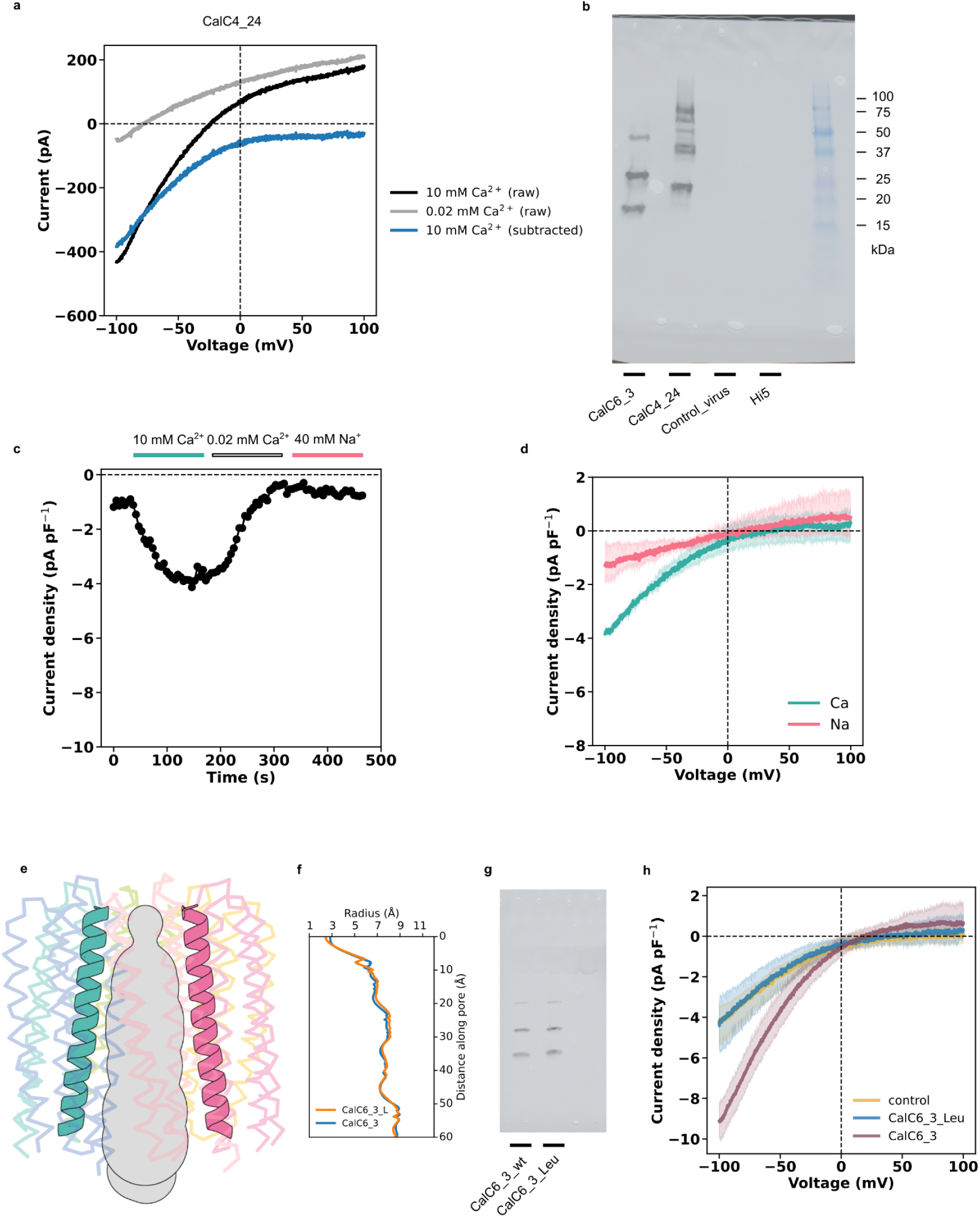
Additional characterizations of designs and negative controls. **a**, An example of I-V curve plot demonstrating the data processing method used to subtract the background leak currents. The black curve represented the raw data obtained from a –100 mV to +100 mV ramp protocol in 10 mM [Ca^2+^] solution. The gray curve represented the raw data obtained using the same ramp protocol in 0.02 mM [Ca^2+^] solution, and was defined as the background. The blue curve was obtained by subtracting the gray curve from the black curve, and was defined as the Ca^2+^ conductance. The same process was used to determine conductances for other ions as well. **b**, Expression of designs in Hi5 cells examined by western blot using an anti-His antibody. The four samples loaded on the gel (lanes indicated by the black bars) were whole-cell lysates of cells expressing CalC6_3 and CalC4_24, cells infected with the control virus (made from an empty pFastBac_Dual vector), and uninfected cells alone. **c,** The time course of currents at –100 mV in 10 mM [Ca^2+^] and 40 mM [Na^+^] extracellular solutions recorded on cells infected with the control virus. **d**, I-V relations obtained from a –100 mV to +100 mV ramp protocol in 10 mM [Ca^2+^] (cyan) and 40 mM [Na^+^] (red) solutions, averaged over three separate cells infected with the control virus. **e,** The side view of the AlphaFold2-predicted model of the CalC6_3-E101L mutant and the calculated ion permeation pathway (using MOLEOnline). **f,** Comparison of the pore radius profiles between the wild type design CalC6_3 (blue) and its E101L mutant (orange). **g,** Western blot analysis of protein expression of the CalC6_3 design and its E101L mutant in Hi5 cells. **h,** I-V relations obtained from a –100 mV to +100 mV ramp protocol in 10 mM [Ca^2+^] on cells expressing the CalC6_3-E101L mutant (blue, n=3), overlaid with the negative control (yellow) and the wild type design CalC6_3 (purple) that are shown in Fig. 3.

**Extended Data Fig. 7:**
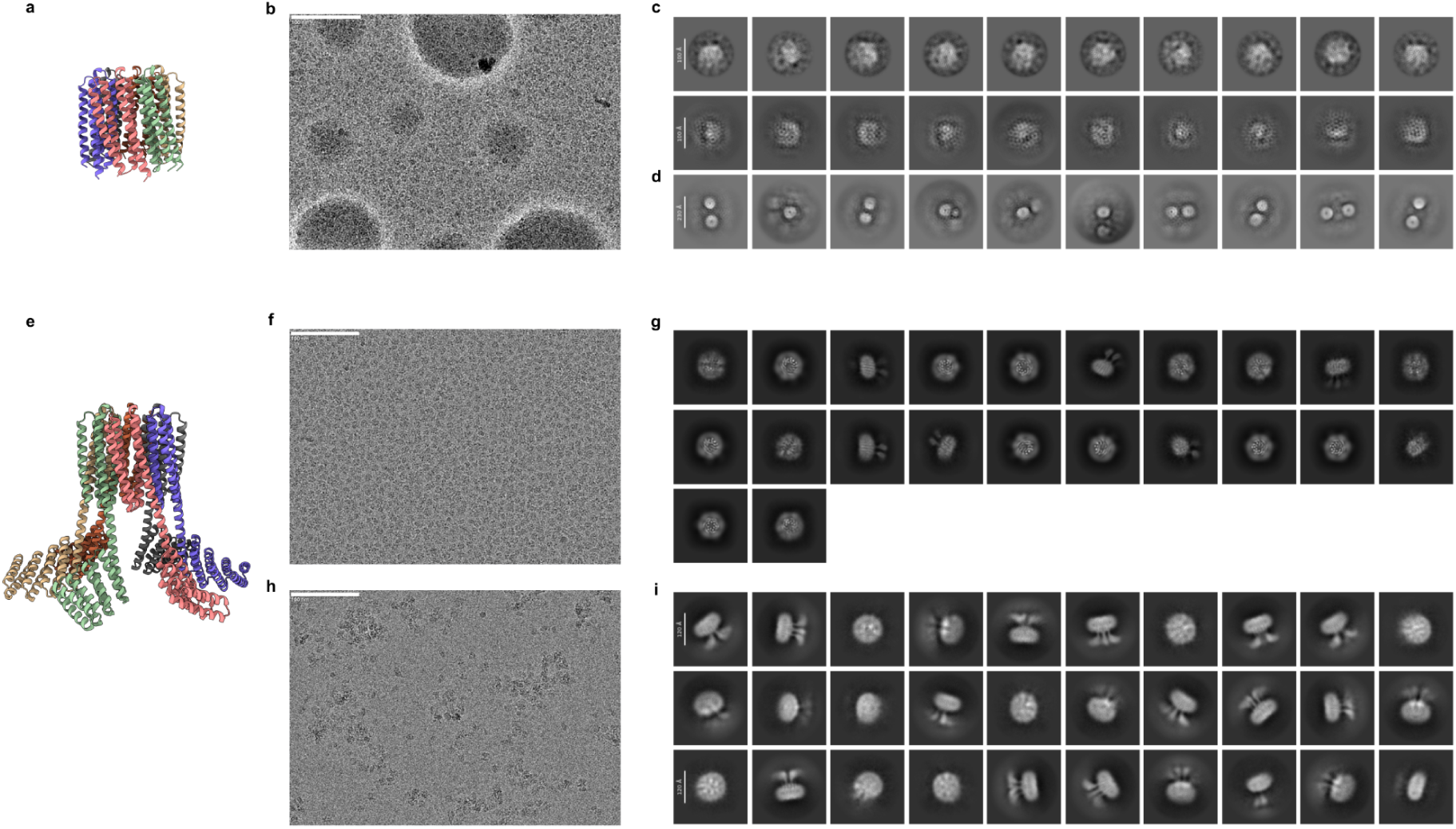
Cryo-EM micrographs and 2D averages for CalC6_3 before and after DHR extensions. **a-d**, Design model **(a)**, a representative cryo-EM micrograph of protein particles on a lacey carbon grid **(b)**, and 2D class averages (**c-d**) from three different 2D classification settings, with an extraction box size of 300 pixels used in **c** and an extraction box size of 680 pixels (pixel size 0.4135 Å) used in **d**. **e-i**, Design model **(e)** and cryo-EM data **(f-i)** for CalC6 with DHR extensions. **f** and **h**, Representative cryo-EM micrographs of protein particles on a thin carbon grid **(f)** and a holey carbon grid **(h)**, respectively. **g** and **i**, 2D class averages from the thin carbon grid **(g)** and from the holey carbon grid **(h)**, respectively. Scale bars in the micrographs, 100 nm. Scale bars in **c**, 10 nm. Scale bar in **d**, 23 nm. Scale bars in **i**, 12 nm.

**Extended Data Fig. 8:**
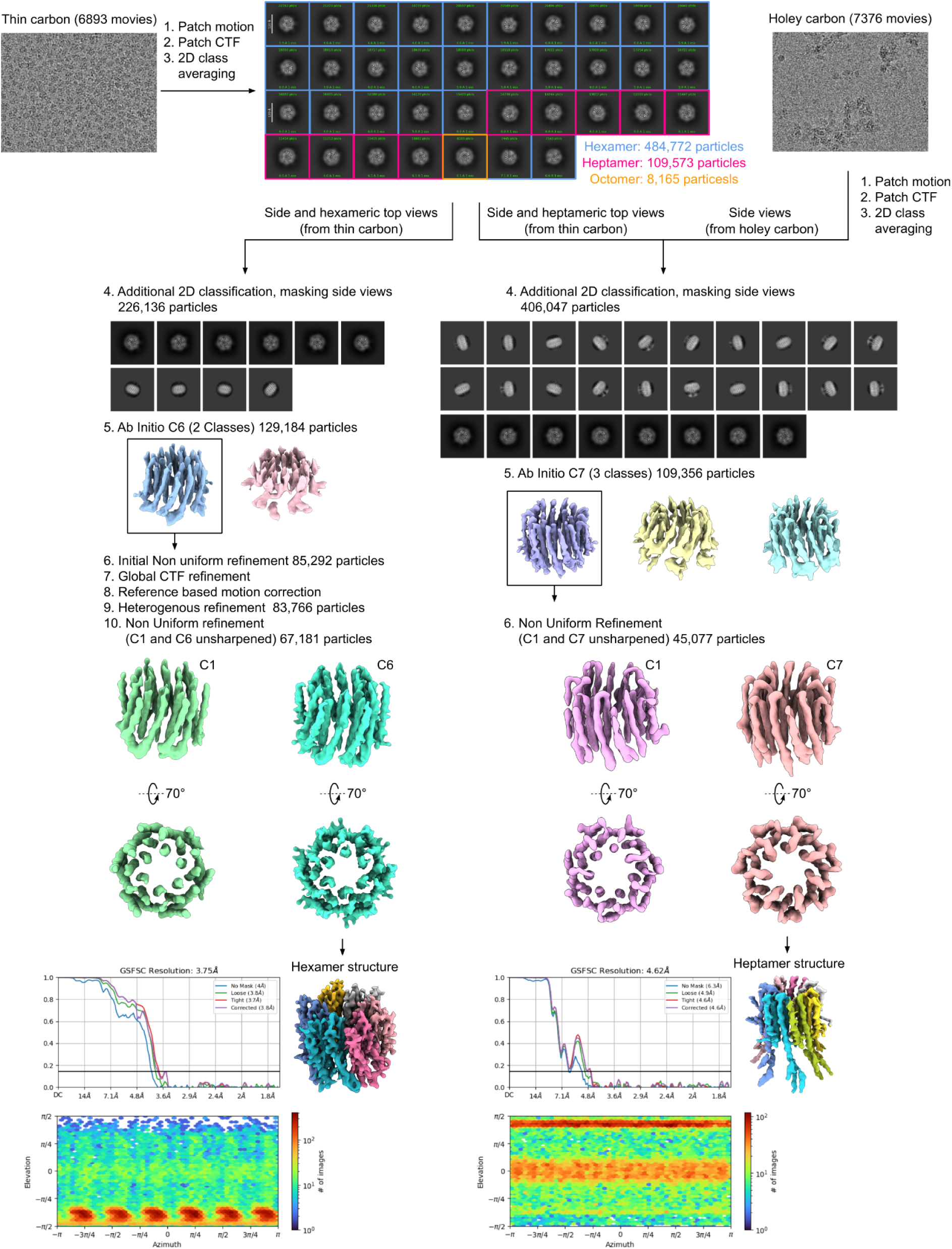
Workflow for cryo-EM data processing of the CalC6_3 design with DHR extensions. The flowchart demonstrated particle picking, classification, reconstruction and refinement processes that enabled determination of both a hexamer and a heptamer structure. All data processing steps were carried out in CryoSPARC v4.4. Detailed processes can be found in Methods (2.11 and 2.12).

**Extended Data Fig. 9:**
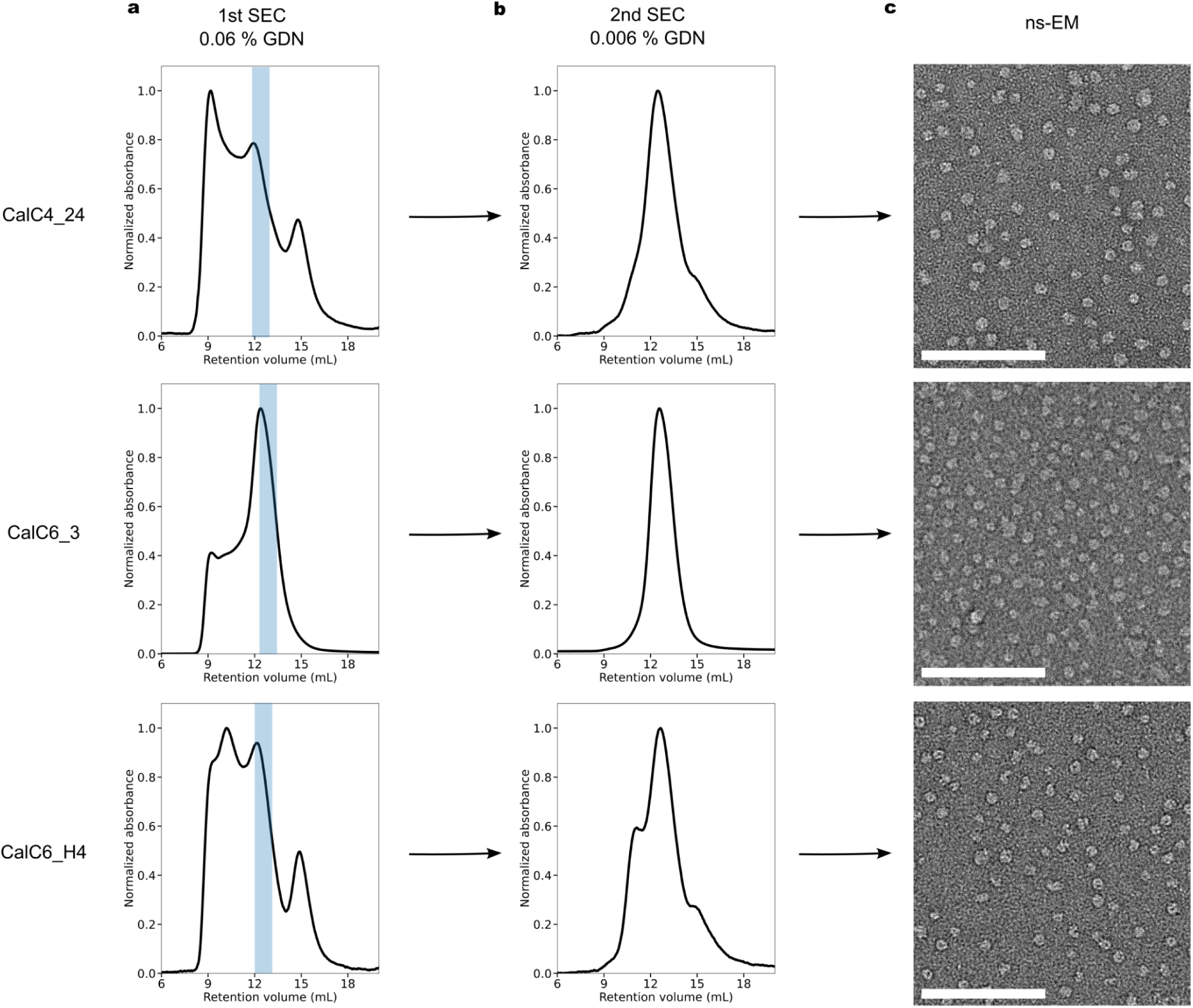
Illustration of the purification strategy for separating the target protein species with the desired oligomeric states from aggregates. The three designs (CalC4_24, CalC6_3 and CalC6_H4), which were previously shown in Fig. 2, were presented here as examples (top, middle and bottom, respectively). From the first SEC run in the buffer containing 0.06% GDN **(a)**, only the latter portions (blue regions) were taken and were subject to the second SEC run in the buffer containing 0.006% GDN **(b,** also shown in **Fig 2d**). Protein samples taken from the more distinct elution peaks were observed to form homogeneous pore-containing particles on ns-EM grids **(c)**. Scale bars, 100 nm.

**Extended Data Table 1:** Cryo-EM data collection and processing parameters for CalC6_3 with DHR extension domains.

|  |  |  |
| --- | --- | --- |
| **Data Collection** |  |  |
| Microscope | Titan Krios |  |
| Voltage (kV) | 300 |  |
| Detector | Gatan K3 |  |
| Energy Filter | Gatan BioQuantum Gif |  |
| Recording mode | Counting |  |
| Magnification | 105,000 X |  |
| Movie micrograph pixel size (Å) | 0.843 |  |
| Dose rate (e<sup>-</sup>/Å<sup>2</sup>/s) | 11.83 |  |
| No. of frames per movie micrograph | 79 |  |
| Frame exposure time (s) | 0.0506 |  |
| Movie micrograph exposure time (s) | 4.0 |  |
| Total dose (e<sup>-</sup>/Å<sup>2</sup>) | 47.32 |  |
| Under focus range (μm) | 0.8-1.8 |  |
| Number of movie micrographs | 14,269 |  |
| **Map Processing** | Hexamer | Heptamer |
| Extraction Box Size (pix) | 340 | 340 |
| Initial particle images (no.) | 886,110 | 886,110 |
| Final particle images (no.) | 67,181 | 45,077 |
| Map resolution (Å) | 3.75 | 4.62 |
| FCS threshold | 0.143 | 0.143 |
| Map resolution range (Å) | 3.38-4.56 | 4.20-8.80 |
| **Refinement** |  |  |
| Initial model used | Design Model | Design Model |
| Map resolution (Å) | 3.75 | 4.62 |
| FCS threshold | 0.143 | 0.143 |
| Model resolution range | 3.38-4.56 | 4.20-6.5 |
| Map sharpening B factor | Na | 343.8 |
| Model composition |  |  |
| Non-hydrogen atoms | 6,215 | 6,069 |
| Protein Residues | 846 | 1,057 |
| Ligands | 0 | 0 |
| *B* factors (Å) |  |  |
| Protein | Na | 249.89 |
| Ligands | Na | – |
| R.M.S. deviations |  |  |
| Bond lengths (Å) | 0.012 | 0.001 |
| Bond angles (°) | 1.795 | 0.369 |
| Validation |  |  |
| MolProbity score | 1.00 | 0.69 |
| Clashscore | 0.78 | 0.57 |
| Rotamer Outliers (%) | 2.13 | 0.00 |
| Ramachandran plot |  |  |
| Favored (%) | 98.32 | 99.33 |
| Allowed (%) | 1.68 | 0.67 |
| Disallowed (%) | 0.00 | 0.00 |

## Notes

### Competing Interest Statement

The authors have declared no competing interest.

## References

1. Magnus, C. J. et al. Ultrapotent chemogenetics for research and potential clinical applications. Science 364, eaav5282 (2019).

2. Jing, M. et al. A genetically encoded fluorescent acetylcholine indicator for in vitro and in vivo studies. Nat. Biotechnol. 36, 726–737 (2018).

3. Gouaux, E. & MacKinnon, R. Principles of Selective Ion Transport in Channels and Pumps. Science 310, 1461–1465 (2005).

4. Yue, L., Navarro, B., Ren, D., Ramos, A. & Clapham, D. E. The Cation Selectivity Filter of the Bacterial Sodium Channel, NaChBac. J. Gen. Physiol. 120, 845–853 (2002).

5. Tang, L. et al. Structural basis for Ca^2+^ selectivity of a voltage-gated calcium channel. Nature 505, 56–61 (2014).

6. Nilius, B. et al. The Single Pore Residue Asp542 Determines Ca^2+^ Permeation and Mg^2+^ Block of the Epithelial Ca^2+^ Channel*. J. Biol. Chem. 276, 1020–1025 (2001).

7. Ellinor, P. T., Yang, J., Sather, W. A., Zhang, J.-F. & Tsien, R. W. Ca^2+^ channel selectivity at a single locus for high-affinity Ca^2+^ interactions. Neuron 15, 1121–1132 (1995).

8. Yang, J., Elllnor, P. T., Sather, W. A., Zhang, J.-F. & Tsien, R. W. Molecular determinants of Ca^2+^ selectivity and ion permeation in L-type Ca^2+^ channels. Nature 366, 158–161 (1993).

9. Meyer, J. O. et al. Disruption of the Key Ca^2+^ Binding Site in the Selectivity Filter of Neuronal Voltage-Gated Calcium Channels Inhibits Channel Trafficking. Cell Rep. 29, 22–33.e5 (2019).

10. Long, S. B., Tao, X., Campbell, E. B. & MacKinnon, R. Atomic structure of a voltage-dependent K+ channel in a lipid membrane-like environment. Nature 450, 376–382 (2007).

11. Derebe, M. G., Zeng, W., Li, Y., Alam, A. & Jiang, Y. Structural studies of ion permeation and Ca^2+^ blockage of a bacterial channel mimicking the cyclic nucleotide-gated channel pore. Proc. Natl. Acad. Sci. 108, 592–597 (2011).

12. Shen, P. S. et al. The Structure of the Polycystic Kidney Disease Channel PKD2 in Lipid Nanodiscs. Cell 167, 763–773.e11 (2016).

13. Magnus, C. J. et al. Chemical and Genetic Engineering of Selective Ion Channel–Ligand Interactions. Science 333, 1292–1296 (2011).

14. Wu, Z. et al. A sensitive GRAB sensor for detecting extracellular ATP in vitro and in vivo. Neuron 110, 770–782.e5 (2022).

15. Joh, N. H. et al. De novo design of a transmembrane Zn^2+^-transporting four-helix bundle. Science 346, 1520–1524 (2014).

16. Mravic, M. et al. Packing of apolar side chains enables accurate design of highly stable membrane proteins. Science 363, 1418–1423 (2019).

17. Mahendran, K. R. et al. A monodisperse transmembrane α-helical peptide barrel. Nat. Chem. 9, 411–419 (2017).

18. Krishnan R, S. et al. Assembly of transmembrane pores from mirror-image peptides. Nat. Commun. 13, 5377 (2022).

19. Lear, J. D., Wasserman, Z. R. & DeGrado, W. F. Synthetic Amphiphilic Peptide Models for Protein Ion Channels. Science (1988) doi:10.1126/science.2453923.

20. Xu, C. et al. Computational design of transmembrane pores. Nature 585, 129–134 (2020).

21. Scott, A. J. et al. Constructing ion channels from water-soluble α-helical barrels. Nat. Chem. 13, 643–650 (2021).

22. Berhanu, S. et al. Sculpting conducting nanopore size and shape through de novo protein design. Science 385, 282–288 (2024).

23. Vorobieva, A. A. et al. De novo design of transmembrane β barrels. Science 371, eabc8182 (2021).

24. Harding, M. M. The geometry of metal–ligand interactions relevant to proteins. Acta Crystallogr. D Biol. Crystallogr. 55, 1432–1443 (1999).

25. Harding, M. M. The geometry of metal–ligand interactions relevant to proteins. II. Angles at the metal atom, additional weak metal–donor interactions. Acta Crystallogr. D Biol. Crystallogr. 56, 857–867 (2000).

26. Elinder, F. & Århem, P. Metal ion effects on ion channel gating. Q. Rev. Biophys. 36, 373–427 (2003).

27. Corry, B., Allen, T. W., Kuyucak, S. & Chung, S.-H. Mechanisms of Permeation and Selectivity in Calcium Channels. Biophys. J. 80, 195–214 (2001).

28. Hess, P. & Tsien, R. W. Mechanism of ion permeation through calcium channels. Nature 309, 453–456 (1984).

29. Sather, W. A. & McCleskey, E. W. Permeation and Selectivity in Calcium Channels. Annu. Rev. Physiol. 65, 133–159 (2003).

30. Almers, W., McCleskey, E. W. & Palade, P. T. A non-selective cation conductance in frog muscle membrane blocked by micromolar external calcium ions. J. Physiol. 353, 565–583 (1984).

31. Cibulsky, S. M. & Sather, W. A. The Eeee Locus Is the Sole High-Affinity Ca^2+^ Binding Structure in the Pore of a Voltage-Gated Ca^2+^ Channel: Block by Ca^2+^ Entering from the Intracellular Pore Entrance. J. Gen. Physiol. 116, 349–362 (2000).

32. Tang, L. et al. Structural basis for inhibition of a voltage-gated Ca^2+^ channel by Ca^2+^ antagonist drugs. Nature 537, 117–121 (2016).

33. Wu, J. et al. Structure of the voltage-gated calcium channel Ca_v_1.1 at 3.6 Å resolution. Nature 537, 191–196 (2016).

34. Saotome, K., Singh, A. K., Yelshanskaya, M. V. & Sobolevsky, A. I. Crystal structure of the epithelial calcium channel TRPV6. Nature 534, 506–511 (2016).

35. Hou, X., Outhwaite, I. R., Pedi, L. & Long, S. B. Cryo-EM structure of the calcium release-activated calcium channel Orai in an open conformation. eLife 9, e62772 (2020).

36. Watson, J. L. et al. De novo design of protein structure and function with RFdiffusion. Nature 620, 1089–1100 (2023).

37. Dauparas, J. et al. Robust deep learning–based protein sequence design using ProteinMPNN. Science 378, 49–56 (2022).

38. Jumper, J. et al. Highly accurate protein structure prediction with AlphaFold. Nature 596, 583–589 (2021).

39. Zhu, W., Shenoy, A., Kundrotas, P. & Elofsson, A. Evaluation of AlphaFold-Multimer prediction on multi-chain protein complexes. Bioinformatics 39, btad424 (2023).

40. Feldman, D. et al. Optical Pooled Screens in Human Cells. Cell 179, 787–799.e17 (2019).

41. Abramson, J. et al. Accurate structure prediction of biomolecular interactions with AlphaFold 3. Nature 630, 493–500 (2024).

42. Vennekens, R. et al. Permeation and Gating Properties of the Novel Epithelial Ca^2+^ Channel*. J. Biol. Chem. 275, 3963–3969 (2000).

43. Hille, B. Ion Channels of Excitable Membranes. (Sinauer Assoc, Sunderland, Mass, 2001).

44. Voets, T. et al. CaT1 and the Calcium Release-activated Calcium Channel Manifest Distinct Pore Properties*. J. Biol. Chem. 276, 47767–47770 (2001).

45. Yue, L., Peng, J.-B., Hediger, M. A. & Clapham, D. E. CaT1 manifests the pore properties of the calcium-release-activated calcium channel. Nature 410, 705–709 (2001).

46. McNally, B. A., Somasundaram, A., Yamashita, M. & Prakriya, M. Gated regulation of CRAC channel ion selectivity by STIM1. Nature 482, 241–245 (2012).

47. Prakriya, M. The molecular physiology of CRAC channels. Immunol. Rev. 231, 88–98 (2009).

48. Hoth, M. & Penner, R. Depletion of intracellular calcium stores activates a calcium current in mast cells. Nature 355, 353–356 (1992).

49. Lansman, J. B., Hess, P. & Tsien, R. W. Blockade of current through single calcium channels by Cd^2+^, Mg^2+^, and Ca^2+^. Voltage and concentration dependence of calcium entry into the pore. J. Gen. Physiol. 88, 321–347 (1986).

50. Bers, D. M., Patton, C. W. & Nuccitelli, R. A Practical Guide to the Preparation of Ca2+ Buffers. in Methods in Cell Biology (ed. Whitaker, M.) vol. 99 1–26 (Academic Press, 2010).

51. Hou, X., Burstein, S. R. & Long, S. B. Structures reveal opening of the store-operated calcium channel Orai. eLife 7, e36758 (2018).

52. Zhang, K., Wu, H., Hoppe, N., Manglik, A. & Cheng, Y. Fusion protein strategies for cryo-EM study of G protein-coupled receptors. Nat. Commun. 13, 4366 (2022).

53. Brunette, T. J. et al. Exploring the repeat protein universe through computational protein design. Nature 528, 580–584 (2015).

54. Drożdżyk, K. et al. Cryo-EM structures and functional properties of CALHM channels of the human placenta. eLife 9, e55853 (2020).

55. Demura, K. et al. Cryo-EM structures of calcium homeostasis modulator channels in diverse oligomeric assemblies. Sci. Adv. 6, eaba8105 (2020).

