## Supplementary information for "Bottom-up design of calcium channels from defined selectivity filter geometry"

6 We would like to dedicate this paper to the memory of William Catterall.

### Methods

#### 1. Computational design methods

#### 1.1. Placement of pore helices

The pore helices can be generated in three steps: 1) placement of $Ca^{2+}$-coordinating residues as the selectivity filter, 2) placement of pore exit residues, and 3) generation of protein backbones to hold these pore-defining residues.

In theory any structural element containing carboxylates and $Ca^{2+}$ ions can be used to define the filter. We extracted the six Glu residues from the open-state structure of Orai channel as the template $Ca^{2+}$-coordinating motif to start from. The coordinates of the motif were changed simultaneously such that the motif was placed in the x-y plane with the $Ca^{2+}$ ion on the z axis using PyMOL. Then five copies of the Glu/Asp residues were deleted, and the x and y coordinates of the remaining Glu/Asp residue were sampled using PyRosetta through rigid-body moves to generate ion-residue pairs with different distances. These ion-residue pairs were symmetrized using the SetupForSymmetry Mover in RosettaScripts with applied C4 or C6 symmetry to generate selectivity filters with different geometric parameters.

The pore exit residues were generated by positioning a set copy of the $Ca^{2+}$-coordinating residues at specific distances (h) along the z axis away from the original x-y plane. This positioning was applied as rigid body transformation in Pyrosetta. The z-distance between the two layers of residues defined the length of the pore. The ideal z-distance was determined to be 43.2 Å in our case mainly based on three considerations: (1) the vertical distance between residue i and i+8 that have the same sidechain orientation (i.e., two consecutive helical turns) is 10.8 Å, (2) the membrane thickness in eukaryotic cells is usually greater than 30 Å[56,57], and (3) we want the pore helices to extend beyond the lipid bilayer, which can reduce the overall hydrophobicity of these helices, and in turn reduce the trend of protein aggregation and non-specific interhelical packing interactions driven by large patches of hydrophobic residues. So the pore length is determined to be 4 x 10.8 Å = 43.2 Å. At this stage the specific amino acid identities of the pore exit residues are inconsequential, as the sequence will be redesigned by ProteinMPNN at a later sequence design step. The x and y coordinates of these pore exit residues were sampled using the rigid body transformation function of Pyrosetta to generate pores with different geometries.

To form the lining of the pore, RFdiffusion was used to symmetrically generate helices that connected the selectivity filter and pore exit residues. The number of the helical linker residues was sampled in order to generate straight alpha helices as the optimal secondary structure. Typically 25-29 residues were sufficient to position the pore entrance and exit residues on a straight helix, depending on the tilt angle of the pore helix relative to the z axis.

#### 1.2. Generation of supporting protein backbones

Once the pore helices were obtained, RFdiffusion was subsequently used to symmetrically generate protein backbones for extending the pore helices into homo-oligomeric subunits with L residues in each monomer. The overall length of the monomer (L) largely depends on the number of times it passes through the membrane and relative distances between neighboring chains. In particular, the number of residues before (at N-terminal of) and after (at C-terminal of) the pore helices were sampled to ensure adequate buttressing of protein backbones within each monomer and extensive interface areas between neighboring subunits. In general, we found that 30-40 residues were sufficient to form a straight single-pass helix, and could be used as the repeating unit to estimate a rough range of the residue numbers needed to generate. The backbone outputs were examined by eye, and those with well-buttressed helices and open pores were selected for the subsequent protein sequence design.

We separated the processes of the generation of pore helices and the scaffolding of pore helices although both steps were done by RFdiffusion. This appeared to be more effective in guiding RFdiffusion to make more compact ion channel-like helical structures than only scaffolding the selectivity filter and pore exit residues. In the latter case, the output backbones mostly contained fiber-like helical coiled-coil assemblies extending far away from the selectivity filter.

#### 1.3. Fine-tuning RFDiffusion on transmembrane proteins

When the diameter of the pore exit is large (> 20 Å between diagonal Ca atoms), the published RFdiffusion model[36] tends to generate protein fragments inside the pore, thus occluding the ion permeation pathway. This is likely due to the globular nature of soluble proteins that constituted the majority of the training dataset for RFdiffusion. To overcome this, we fine-tuned RFdiffusion on a dataset containing 6,392 transmembrane proteins from the OPM (https://opm.phar.umich.edu/) database comprising all beta-barrel and alpha-helical proteins as of April 2023 to generate backbones with higher chances of maintaining a pore. Specifically, we fine-tuned the version of RFdiffusion trained with (partial) secondary structure and block-adjacency information provided, and continued to provide such information during fine-tuning, such that at design time one can specify the desired protein topology. We fine-tuned RFdiffusion in precisely the manner described in Watson et al., 2022, and trained for 100 epochs on this transmembrane dataset.

1.4. Sequence design on protein backbones

The selected protein backbones of the channels were sequence-designed with ProteinMPNN. The --tied_positions argument was used to restrict sequences to be identical on each monomer. The residues forming the selectivity filter were kept undesigned using the --fixed_positions argument. Position-specific amino acid constraints were applied using the --omit_AAs argument. The pore-lining residues were selected based on the distance between the Cα atom of the residue and the central z axis (the square root of the sum of the squares of x and y coordinates of that CA atom in the PDB file). Charged and bulky aromatic amino acids (tyrosine and tryptophan) were excluded from the selected pore residues to prevent interference with ion permeation through the pore, particularly through sites that might form narrow constrictions. Residues that were defined as being embedded in the lipid bilayer were selected based on their relative vertical (z) distances to the selectivity filter. The selectivity filter residues were arbitrarily designated as the initial point of the lipid bilayer on the extracellular side. Residues that are within a certain range on the z-distance (e.g., 30 Å) were all defined to be embedded in the lipid bilayer. The lipid-facing residues were selected combinatorially by (1) the Rosetta LayerSelector using the sidechain neighbor algorithm, (2) not being among the pore-lining residues, and (3) being among the lipid-embedded residues. Tyrosine (Tyr) and tryptophan (Trp) residues were placed at the first and final residues of the surface residues on a consecutive helix, respectively, interfacing the aqueous phase and lipid phase.

It is optional to use the ProteinMPNN model trained with extra input per residues specifying buried and interface residues[58] using the argument --model_type "per_residue_label_membrane_mpnn". In this case the lipid-facing residues were parsed into ProteinMPNN using the --transmembrane_buried argument. Other constraints were applied in the same way as described above.

1.5. Structure prediction on designed sequences

AlphaFold2 was primarily used to judge if the designed sequences would fold and assemble into homo-oligomers as designed. Model 4 seemed to be most predictive of the designed alpha-helical transmembrane proteins and was the only model used. Designs were first filtered by confidence scores (typically pTM > 0.85 and mean PAE scores < 10 for all residue pairs corresponding to different chains) and then manually inspected to exclude designs with clusters of hydrophobic residues outside the defined lipid-embedding regions.

For estimating ion binding at the selectivity filters, AlphaFold3 was used to predict the formation of protein-metal complexes. Note that there could be clusters of negatively charged residues on the surface of the designed proteins forming potential ion binding sites. Therefore, multiple copies of ions were often included as the input rather than a single ion.

#### 1.6. Resampling and optimization of protein backbones

In some cases, very few designed sequences were predicted to fold with high confidences for some specific protein backbones with very tilted pore geometries. We hypothesized that the low in silico success rate was due to non-ideality of the protein backbones, which resulted in a low “designability” by ProteinMPNN. To refine the protein backbones, we applied partial noising and denoising processes using RFdiffusion partial diffusion to sample for more reasonable structures around the original backbone. This resampling process was not applied on the pore helices to maintain the pore geometry. The newly generated backbones were then subjected to another round of backbone selection, sequence design, and structural prediction as described above.

#### 1.7. Connecting DHR to designed oligomeric channel for Cryo-EM analysis

To connect DHR proteins as the soluble domain to CalC6_3 for structural determination, the main objective is to avoid potential steric clashes between the subunits after addition of multiple copies of DHR proteins. This was achieved by precisely adjusting the relative position of the DHR proteins to CalC6_3, and this positioning operation was primarily done in PyMOL. The N-termini of experimentally-validated DHR proteins with different twisted geometries were aligned to the C-terminus of the CalC6-3 monomer and were oriented to face away from the pore. Then the DHRs were moved in the direction parallel to the aligned helix away from CalC6_3 to a point where the DHRs were unlikely to clash with the neighboring subunits. RFdiffusion was subsequently used to generate straight helices to fuse DHRs to the CalC6_3 monomer. The linked structure was symmetrized using the Rosetta SetupForSymmetry Mover by generating five extra copies. The residues that might potentially interact with neighboring subunits were manually selected and redesigned by ProteinMPNN as homo-hexamer. The sequence of the redesigned monomer was predicted by AlphaFold2 (the whole complex was too large for AlphaFold2 to predict). Designs with lowest Cα RMSDs to the CalC6_3 model in regard to the channel region were selected for experimental characterization.

### 2. Experimental methods

2.1 Construction of synthetic genes

The amino acid sequences of the designs were reverse translated to DNA sequences using DNAworks (Hoover, D. and Lubkowski, J., 2002, https://github.com/davidhoover/DNAWorks). Synthetic genes encoding the designs were ordered from Genscript Inc. (Piscataway, N.J., USA) or Integrated DNA Technologies, Inc. (IDT, Coralville, Iowa, USA). The genes encoding the designs for protein expression and purification were cloned into pET29b+ vector between NdeI and XhoI sites with a C-terminal hexahistidine tag. The genes encoding the designs for expression in HEK293T cells were cloned into the LentiGuide-BC-EF1a vector between XbaI and EcoRI sites with a C-terminal FLAG tag (DYKDDDDK). The genes encoding the designs for expression in insect cells were cloned into the MCS-1 of pFastBac-Dual vector between BamHI and XbaI sites with a C-terminal hexahistidine tag.

2.2. Expression of designed channels in HEK cells

HEK293T cells were maintained at 37 °C with 5% $CO_2$ and cultured in Dulbecco's Modified Eagle's Medium (DMEM, Gibco) supplemented with 10% FetalClone II serum (FC-II, Cytiva HyClone) and 1% penicillin-streptomycin (Gibco). To favor a more homogeneous expression level for more consistent flux assay results, we expressed the designed channels in HEK293T cells by lentivirus transduction based on a published protocol with some minor modifications[40].

To prepare the lentivirus, cells were transfected with the design plasmid, the envelope plasmid pMD2.G (Addgene #12259) and the packaging plasmid psPAX2 (Addgene #12260) at a 3:4:5 mass ratio using Lipofectamine 3000 (Thermo Fisher L300015). Media was exchanged after 4 hours. At 24 hours post-transfection, caffeine was supplemented to a final concentration of 2 mM to generate virus with higher titer. The virus-containing supernatant was harvested 72 hours after transfection and filtered through 0.45 μm surfactant-free cellulose acetate (SFCA) filters (Corning 431220). The prepared viruses were used immediately or stored at -80 °C.

For lentiviral transduction, cells were dissociated from the culture flask using 0.05% trypsin-EDTA (Gibco) and resuspended in DMEM-10% FC-II medium supplemented with polybrene (8 μg/mL, Santa Cruz Biotechnology) at a cell density of 1,000,000 cells/mL. Viral supernatant was added to this cell suspension (typically 200 μL viral supernatant was used to transduce 1 mL of the above cell suspension). The cells-virus mixture was centrifuged at 1000 g for 2 hours at 33°C. Media was exchanged 24 hours after infection.

2.3. Flux assay of designed channels in HEK cells

At 48 hours post-infection, transduced cells were seeded into glass-bottom 96-well plates (Cellvis P96-1.5H-N) pretreated with 0.05% poly-D-lysine hydrobromide (Sigma, dissolved in sterile-filtered Milli-Q water) at a density of 60,000 cells per well. A $Ca^{2+}$-free HEPES-buffered solution (20 mM HEPES pH 7.4, 150 mM NaCl, 5 mM KCl, 2 mM $MgCl_2$, 10 mM glucose, osmolarity adjusted to 330 mmol/kg using sucrose) was used as the assay buffer. To prepare the dye-loading solution, a vial of Fura-2 AM dye (Invitrogen F1221, 50 μg) was thawed, resuspended in 10 μL DMSO, and added to the assay buffer supplemented with Powerload (Invitrogen P10020) and probenecid (Invitrogen P36400) to a final dye concentration of 5 μM. Cell culture media was replaced with this dye-loading solution and incubated for 1 hour at room temperature. After the dye loading step, the cells were washed with the assay buffer twice, incubated in the assay buffer, and measured for fluorescence at 510 nm with excitations at 340 nm and 380 nm (F340/F380) using the Neo2 plate reader. The measurement was performed for 15 minutes to obtain the baseline fluorescence. Subsequently, $Ba^{2+}$ was added to the cells to a final concentration of 2 mM and the fluorescence was measured for 15 minutes to obtain the change of fluorescence ratio over time. Data were exported as Excel tables, processed and plotted using Python.

### 2.4. Protein production in E.coli

The designs in pET29b+ vector were transformed into E. coli expression strain BL21(DE3*) (New England Biolabs, MA, USA). The transformed cells were grown in 50 mL Terrific Broth II (TB-II, MP Biomedicals) medium with a final concentration of 50 μg/ml kanamycin in 250 mL baffled flasks overnight. The 50 mL cultures were inoculated into 500 mL TB-II medium in 2 L baffled flasks and incubated at 37°C. After 3 hours, isopropyl β-D-1-thiogalactopyranoside (IPTG) was added to a final concentration of 0.5 mM. The cultures were incubated at 18°C following the addition of IPTG and were harvested after 3 hours by centrifuging at 4000 g for 10 minutes at 12°C. The cell pellets can be stored at -80°C.

### 2.5. Buffer recipe for protein purification

Lysis buffer: 20 mM Tris-HCl, 150 mM NaCl, pH 8.0, supplemented with Pierce Protease Inhibitor Tablets (Thermo Scientific A32963, 1 tablet per 100 mL solution)

Solubilization buffer: 20 mM Tris-HCl, 150 mM NaCl, pH 8.0, supplemented with 2% w/v n-Decyl-β-D-Maltopyranoside (DM, Anatrace D322)

Wash buffer 1: 50 mM Tris-HCl, 300 mM NaCl, 30 mM imidazole, pH 8.0, supplemented with 0.06% w/v glyco-diosgenin (GDN, Anatrace GDN101)

Wash buffer 2: 50 mM Tris-HCl, 1 M NaCl, 30 mM imidazole, pH 8.0, 0.06% GDN

Wash buffer 3: 50 mM Tris-HCl, 300 mM NaCl, 60 mM imidazole, pH 8.0, 0.06% GDN

Elution buffer: 50 mM Tris-HCl, 300 mM NaCl, 500 mM imidazole, pH 8.0, 0.06% GDN

SEC buffer 1: 20 mM Tris-HCl, 150 mM NaCl, pH 8.0, 0.06% GDN

SEC buffer 2: 20 mM Tris-HCl, 150 mM NaCl, pH 8.0, 0.006% GDN

2.6. Purification of designed calcium channels

The cell pellets were resuspended in 30 mL lysis buffer. The suspension was placed on ice and lysed by sonication (QSonica Sonicators, CT, USA) for 4 minutes (15 seconds on/15 seconds off, 8 minutes total run time) at 65% power with ¾'' Dia replaceable tips. The lysates were centrifuged at 10000 g for 15 min to remove the cell debris. The supernatant was collected and ultracentrifuged at 170,000 g for 1 hour at 12°C using Beckman Optima XE-90 Ultracentrifuge to pellet down the membrane fraction. 8 mL solubilization buffer was added to the pellet and incubated at 4°C overnight on a rocker. The homogenate was centrifuged at 30,000 g for 30 minutes to pellet down the materials that could not be solubilized by DM. The supernatant was collected and applied to chromatography columns (BIO-RAD Econo-Pac gravity flow columns) containing Ni-NTA resin (Qiagen, MA, USA). The resin was washed with 5x column volume (CV) of wash buffer 1, 5x CV of wash buffer 2 and 3x CV of wash buffer 3. The proteins were eluted with 4x CV of elution buffer, concentrated in 100 kDa molecular weight cutoff spin concentrator (Millipore), and further purified by size exclusion chromatography (SEC) in SEC buffer 1. Normally a Superdex 200 Increase 10/300 GL column was used. For designed proteins connected with DHR, a Superose 6 increase 10/300 column was used. Both types of columns were from Cytiva, MA, USA. The target elution volume containing the desired oligomeric states of the designs was compared with validated de novo designed transmembrane proteins with similar molecular weight (for example, TMH4C4, which approximated a molecular weight of around 100 kDa, and TMHC6, which approximated a molecular weight of around 60 kDa).

In many cases, there seemed to be an evident aggregation peak when samples were applied on the SEC column after Ni-NTA purification (Extended Data Fig. 9). This is a well-known issue for the purification of membrane proteins, whereby the behavior of membrane proteins in vitro is largely dependent on the detergent or lipids utilized. We took a similar purification strategy as for purifying mammalian voltage-gated sodium channels[59–63]: we took the latter portion of the elution peak containing the proteins that were estimated to be in the desired oligomeric states, and subjected this portion to the a second SEC run in a buffer with ten 10 times less detergent (SEC buffer 2). The second

SEC yielded a more distinct peak corresponding to the protein in the desired oligomeric states, which enabled us to separate the species of interest. 0.006 % GDN was used in the final buffer, as this consistently gave the optimal EM results.

### 2.7. Circular dichroism (CD) measurements

CD spectra were measured on J-1500 Circular Dichroism Spectrophotometer (Jasco). Proteins were prepared at ~0.2 mg/mL in SEC buffer 2. Wavelength-scan spectra were recorded from 190 nm to 260 nm (0.1 nm increments). Temperature melting experiments were conducted from 25 °C to 95 °C (heating rate 1 °C/min), with wavelength-scan spectra recorded at every 10 °C increment. The full spectra were measured again after the samples were cooled down to 25 °C.

### 2.8. Negative-stain electron microscopy

Proteins were prepared at ~0.05 mg/mL in SEC buffer 2 for ns-EM. 6 µL of the protein sample was applied on glow discharged, carbon-coated 400-mesh copper grids (Electron Microscopy Sciences). The samples were allowed to adhere to the grids for 1 min before being wicked away. Then each grid was stained with 3 µL of 2% uranyl formate for four times, each time 20 seconds. The grids were air-dried and were imaged on a FEI Talos L120C TEM (FEI Thermo Scientific, Hillsboro, OR) equipped with a Ceta 4K CCD camera at a magnification of 73,000x at 120 kV. Micrographs collection was automated using the EPU software (FEI Thermo Scientific). Collected datasets were imported into CryoSPARC software (v4.6.0) for 2D class averaging. Micrographs were motion-corrected and CTFs were estimated using Patch CTF estimation function. For each dataset, ~200 particles were selected using the manual particle picker function to generate initial 2D class averages using the 2D classification program in CryoSPARC. The 2D class averages were used as templates for automated particle picking using the template particle picking function after which 50 classes of 2D class averaging were generated for analysis.

### 2.9. Cryo-EM Sample preparation

For CalC6_3, 3 µL of protein at ~ 1 mg/mL in SEC buffer 2 (20 mM Tris-HCl pH 8, 150 mM NaCl, 0.006% GDN) was applied to glow-discharged (25 seconds at 15 mA) 3 nm lacey carbon Cu 400 mesh grids (Electron Microscopy Sciences).

For CalC6_3 with DHR extensions, protein was prepared at ~1 mg/mL in SEC buffer 2 (20 mM Tris-HCl pH 8, 150 mM NaCl, 0.006% GDN). 2 µL of protein sample was applied to glow-discharged (25 seconds at 15 mA) C-flat R 2.0/2.0 300 mesh holey carbon grids

(Electron Microscopy Sciences), and 3 µL of protein sample was applied to glow-discharged (16 seconds at 5 mA) Quantifoil 2 nm thin carbon R 2.0/2.0 300 mesh grids.

Vitrification was performed using a Mark IV Vitrobot at 22 °C and 100% humidity for all grids. Blotting was done before immediately plunge-frozen into liquid ethane. A blot time of 0.5 second, a blot force of 0 and a wait time of 7.5 second were used for lacey carbon grids and Quantifoil thin carbon grids. A blot time of 6.5 second, a blot force of 0, and a wait time of 7.5 second were used for holey carbon grids. The grids were then clipped and stored in liquid nitrogen for collection on the Titan Krios.

### 2.10. Cryo-EM data collection

Samples were collected automatically using SerialEM[64], used to control a ThermoFisher Titan Krios 300 kV microscope equipped with a K3 Summit direct electron detector[65] and BioQuantum Gif energy filter. For CalC6_3 with DHR extensions (thin carbon and holey carbon grids) the microscope was operated in counting mode. For CalC6_3 (lacey carbon grids) super resolution mode was used. Data were collected with random defocus ranges spanning between −0.8 and −1.8 µm using beam-image shift five shots per hole and nine holes per stage move were collected for CalC6_3 with DHR extensions (thin and holey carbon grids) and 27 shots per stage movement (3x9) for CalC6_3. Altogether 6,893, 7,376, and 4,644 movies with pixel sizes of 0.843, 0.843, and 0.4135 and doses of 47.32, 47.32, and 60 e-/Å$^2$ were recorded for CalC6_3 with DHR extensions on thin carbon grid, holey carbon grid, and CalC6_3 on a lacey carbon grid, respectively.

### 2.11. Cryo-EM data processing

All data processing was carried out in CryoSPARC v4.4[66]. For all movies Patch Motion then Patch CTF data processing was carried out as an initial processing step. Custom settings are mentioned for refinement jobs in the protocol, otherwise default settings were used.

For CalC6_3: Curate Exposures was used to remove 1,078 micrographs, leaving 3,566 good micrographs. Blob Picking was used with a minimum particle diameter of 80 Å and a maximum particle diameter of 120 Å. An extraction box size of 300 pixels, and 680 pixels fourier cropped to 340 pixels was used. Following an Inspect Picks job 1,311,993, and 1,094,418 particles were extracted for 300 and 680 pixel sizes, respectively. Extensive 2D classification was performed on the 300 pixel size particles, however particles could not be effectively classified. The larger extraction box size of 680 yielded some low resolution top views showing the pore (Extended Data Fig. 7a-d). Further 2D classification and possible 3D refinement was abandoned in favor of the CalC6_3 with DHR extensions.

For CalC6_3 with DHR extensions: Curate Exposures was used to remove 172 and 1,217 bad micrographs, leaving 6,721 and 6,161 micrographs for thin and holey carbon grids respectively. Blob picking was used with a minimum particle diameter of 80 Å and maximum particle diameter of 160 Å. Particles were extracted at 340 pixels. Following an Inspect Picks job 8,115,000 and 7,293,711 particles were extracted for thin and holey carbon respectively. For thin carbon particles an initial round of 2D classification, 150 classes with a circular mask of 220 Å was used. From this 886,110 particles representing top and side views were further 2D classified with: 150 classes, batchsize per class of 400, 50 online-EM iterations, min over scale after first iteration set to true, with 3 final iterations using all particles. For holey carbon particles an initial round of 2D classification with: 400 classes, and number of final iterations set to 5 was used. From this 185,009 particles representing top and side views were further 2D classified: 50 classes, and 5 final iterations with all particles. Top view classification was abandoned for holey carbon as no secondary structure could be resolved. From the holey carbon 142,660 particles were identified as side views by the visible DHR extensions. (Extended Data Fig. 7e-i)

For the hexamer structure only the thin carbon data was used. 115,880 particles of top/tilted views were selected from the previous 2D classification job. 110,256 particles with side views were further 2D classified separately using: A maximum resolution and a maximum alignment resolution of 3 Å, circular mask diameter and circular mask diameter outer set to 100Å and 120Å respectively (To mask out the flexible DHR extension), a batch size per class of 400, and number of online-EM iterations set to 40 was used. Of these true side views 13,304 particles were selected where clear secondary structure was visible. These final 129,184 particles with side and top/tilted views were further classified by generating 2 *ab initio* classes with: C6 symmetry, initial lowpass resolution set to 15 Å, GSFSC split resolution set to 10 Å, and optimize per-particle defocus set to true. The best *ab initio* volume and 85,292 particles were used to generate a 4.19 Å volume using Non Uniform refinement with: C6 symmetry, initial lowpass resolution set to 15 Å, GSFSC split resolution set to 10 Å, and optimize per-particle defocus set to true. To correct for any aberrations introduced from the beam-image shift collection strategy, Exposure Group Utilities was used to split the data into 101 exposure groups, using Global CTF Refinement with settings: Fit spherical aberration set to true, fit tetrafoil set to true. The subsequent Non Uniform refinement job improved the resolution of the map to 3.79 Å using: C6 symmetry, initial lowpass resolution set to 15 Å, GSFSC split resolution set to 10 Å, and optimize per-particle defocus set to true. This 3.79 Å map was used as an input for Reference Based Motion Correction, followed by another Non Uniform Refinement which further improved the map resolution to 3.60 Å using the following custom settings: symmetry C6, number of extra final passes 5, initial lowpass resolution set to 15 Å,

GSFSC split resolution set to 10 Å, and optimize per-particle defocus set to true. Using the remaining 83,766 particles and the 3.60 Å map as the input, a round of heterogeneous refinement was performed in C1 with 8 volumes. The particles from the 4 best volumes with clearly defined secondary structure were kept. 67,181 particles were used to generate the final volumes with No Uniform Refinement in C1 and C6 symmetry with: initial lowpass resolution set to 15 Å, GSFSC split resolution set to 10 Å, and optimize per-particle defocus set to true. Final resolutions were 3.75 Å and 3.98 Å for C6 and C1 maps respectively. While a slight drop in resolution was observed the map quality was much higher when visually inspected. The final C6 map used for model building was sharpened with DeepEMhancer[67]. Final maps are deposited in the EMDB under EMD-47340.

For the heptamer structure the thin and holey carbon data was used, utilizing top and tilted views from the thin carbon and side views from both the thin and holey carbon data. From the second 2D classification of the particles on the thin carbon grid, 8 classes of 89,183 particles were selected with top and tilted views that appeared to be larger than hexameric by visual inspection. 110,256 particle side views from the thin carbon grid and 142,660 particle side views from the holey carbon grid were combined and an initial 2D classification was performed with settings: maximum resolution and maximum alignment resolution of 3 Å, circular mask diameter and circular mask diameter outer of 130 Å and 150 Å respectively, batch size of 400, force max over poses/shifts set to false, number of online-EM interactions set to 40. Subsequent 2D class averages were removed that appeared hexameric, judging from the side views used in the hexamer structure. 66,778 particles were further 2D classified using: maximum resolution and maximum alignment resolution of 3 Å, circular mask diameter and circular mask diameter outer of 100 Å and 120 Å respectively, batch size of 400, force max over poses/shifts set to false, number of online-EM interactions set to 40. 46,550 particles were selected where any secondary structure was visible and an additional 2D classification was performed using: maximum resolution and maximum alignment resolution of 3 Å, circular mask diameter and circular mask diameter outer of 100 Å and 120 Å respectively, batch size of 400, force max over poses/shifts set to false, number of online-EM interactions set to 40. 20,173 particles with clear secondary structure were selected. Using the 89,183 particles of heptameric top/tilted views and the 20,173 particles from extensive 2D classification of side views additional sorting was done with an *Ab Initio* job using: 3 *ab initio* classes, C7 symmetry, maximum resolution of 6 Å, Initial resolution of 25 Å. The best *ab initio* volume was selected containing 45,077 particles, and used for a Non-Uniform Refinement job both in C7 and C1 symmetry with: number of extra final passes set to 5, initial lowpass resolution set to 15 Å, GSFSC split resolution set to 10 Å, and optimize per-particle defocus set to true. For the C7 and C1 map a resolution of 4.62 Å and 5.13 Å was achieved respectively. The final C7 map used for model building was sharpened with DeepEMhancer. Final maps are deposited in the EMDB under EMD-47356.

2.12. Model Refinement

For all structures, the design models were used as an initial reference for building the final cryo-EM structures. For the hexamer the design model of the CalC6_3 was used as an initial reference model. For the heptamer, CalC6_3 with DHR extension was used as a starting model, as more of the DHR extension was visible in this map. A single chain for the heptamer was isolated in PyMOL and 7 copies of this chain were fitted to the map in UCSF Chimera[68] to reform the channel. Hexamer and heptamer were refined using several rounds of relaxation and minimization, performed on the complete structures, which were manually inspected for errors each time using Isolde[69], Coot[70,71], and Phenix[72] real-space refinement. The final model quality was analyzed using MolProbity[73]. Figures were generated using UCSF ChimeraX[74]. The final structures were deposited in the Protein Data Bank (PDB) under 9DZW, 9E0H, for the hexamer and heptamer respectively.

2.13. Electrophysiology

We use the Bac-to-Bac system (Gibco) to express the designed channels in *Trichoplusia ni* insect cells (Hi5) for whole-cell patch-clamp experiments. The main advantage of this expression system is that cells expressing the designed proteins can be readily identified based on the morphological changes of Hi5 cells after being infected (characteristic features include an expansion of cell nuclei and an increase in cell granularity).

2.13.1 Insect cell culture

SF9 cells and Hi5 cells were maintained at 27°C and cultured in Grace's Insect Medium supplemented with 10 % fetal bovine serum (Avantor Seradigm 89510-186) and 1% penicillin-streptomycin-glutamine (Cytiva Hyclone). This medium was referred to as "supplemented medium" in the following sections.

2.13.2. Baculovirus generation

The preparation of bacmid and baculovirus can also follow Invitrogen's manual of Bac-to-Bac Baculovirus Expression System available online. We used a protocol that has been developed in the lab and previously used in published research. Designs in pFastBac-Dual vector were transformed into MAX Efficiency™ DH10Bac Competent Cells (Gibco). The transformed E. coli cells were then plated on LB agar plates supplemented with 50 µg/mL kanamycin, 7 µg/mL gentamicin, 10 µg/mL tetracycline, 100 µg/mL Bluo-Gal and

40 μg/mL IPTG and incubated at 37 °C for 48 hours. White colony was picked and transferred to 2 mL LB media supplemented with 50 μg/mL kanamycin, 7 μg/mL gentamicin and 10 μg/mL tetracycline to grow overnight. The cells were harvested by centrifugation at 14,000 g for 2 minutes and resuspended in 300 μL Solution 1 (15 mM Tris-HCl pH 8.0, 10 mM EDTA, 100 μg/ml RNaseA). 300 μL Solution 2 (0.2 M NaOH, 1% SDS) was added to this suspension, followed by gentle inversions several times and incubation at room temperature for 5 minutes to form clear lysates. 300 μL Solution 3 (potassium acetate, pH 5.5) was added to the lysate, followed by incubation on ice for 5 minutes. Cell debris was removed by centrifugation at 14,000 g for 10 minutes. 700 μL supernatant was mixed with 800 μL isopropanol and incubated on ice for 10 minutes. The mixture was centrifuged at 14,000 g for 10 minutes to pellet down the bacmid DNA. After the supernatant was discarded, the pellet was washed with 500 μL 70% ethanol, air dried at room temperature, and resuspended in 40 μL d$H_2O$.

SF9 cells were seeded into 6-well tissue culture dishes at 50% confluency. Grace's Insect Medium without serum or antibiotics (unsupplemented media) was used for transfection at this step. 10 μg bacmid DNA was added into 100 μL unsupplemented medium. 6 μL of Cellfectin II Reagent (Gibco) was diluted by 100 μL of unsupplemented medium and mixed with the bacmid DNA. The mixture was incubated at room temperature for 30 minutes. Pre-seeded SF9 cells were washed once with 2 mL unsupplemented medium. 800 μL of unsupplemented medium was added to the DNA-Cellfectin mixture and added to the cells. Media was exchanged to supplemented medium after incubation at 27°C for 5 hours. The virus-containing supernatant was harvested 72 hours after transfection and was centrifuged at 500 g for 5 minutes to remove cell debris.

The virus (P1) was amplified once to generate a stable stock by addition to the SF9 cells cultured in the 150-mm tissue culture dish (at ~70% confluency). From the culture dish medium was aspirated, and virus suspension was gently added to the cells. The tissue culture dish was put on a rocker at room temperature for 1 hour, and incubated at 27 °C after addition of 25 mL medium. 48 hours after infection the virus-containing supernatant was harvested by centrifugation at 500 g for minutes and was stored at 4 °C (P2 virus).

2.13.3. Western blot analysis

Western blot was used to confirm the expressions of designed channels. Hi5 cells were seeded into a 6-well tissue culture dish at 50% confluency. To infect the cells, medium was aspirated, and 5 μL of the P2 virus was added to the cells. The dish was put on a rocker at room temperature for 1 hour, and incubated at 27 °C after addition of 2 mL medium. Cells were harvested 48 hours after transfection, washed with 1 mL TBS buffer (20 mM Tris-HCl pH 8, 150 mM NaCl) supplemented with pierce protease inhibitor, and resuspended in 60 μL TBS buffer. 5 μL of the cell suspension was added to 50 μL of the

2x Laemmli Sample Buffer (BIO-RAD) supplemented with 2-mercaptoethanol and loaded on gel. Proteins were transferred to the nitrocellulose membrane using the BIO-RAD Trans-Blot Turbo Transfer system. The membrane was incubated with 5% blotting grade blocker nonfat milk (BIO-RAD) in TBST (20 mM Tris-HCl, pH 8, 150 mM NaCl, 0.05% Tween 20) at 4°C overnight. On the following day the membrane was washed three times with TBST and incubated with anti-His tag antibody conjugated with Horseradish Peroxidase (Jackson ImmunoResearch 300-035-240) in TBST (1:5000 dilution) for 1 hour. After antibody binding, the membrane was washed four times with TBST. Chemiluminescence was activated by BIO-RAD Clarity Western ECL Substrate and imaged using LI-COR Odyssey M Imager.

#### 2.13.4. Solution recipe for electrophysiology

All solutions used for electrophysiology were prepared from the stock solutions below:

NMDG-$MeSO_3$ stock solution: 1.4 M NMDG, 200 mM HEPES, pH adjusted to 7.2 by methanesulfonic acid (~1.3 M)

NMDG-Cl stock solution: 1.4 M NMDG, 200 mM HEPES, pH adjusted to 7.2 by HCl (estimated $Cl^-$ concentration of 1.3 M based on the added HCl volume)

$NaMeSO_3$ stock solution: 1.4 M NaOH, 200 mM HEPES, pH adjusted to 7.2 by methanesulfonic acid (~1.3 M)

Stock solutions of divalent ions ($Mg^{2+}$, $Ca^{2+}$, and $Sr^{2+}$) were prepared by mixing the corresponding hydroxides with $MeSO_3$ to a final concentration of 100 mM for the metal ions. A little excess of $MeSO_3$ was usually required to ensure complete dissolving of the metal hydroxides to a final pH of 3~5.

EGTA stock solution: 100 mM EGTA, 200 mM NMDG.

Extracellular solution: 140 mM NMDG, ~130 mM $MeSO_3$, 20 mM HEPES, ~4 mM $Cl^-$, x mM metal ions of interest, pH 7.2, osmolarity adjusted to 355 mmol/kg using sucrose.

Intracellular solution: 160 mM NMDG, ~110 mM $MeSO_3$, 20 mM EGTA, 20 mM HEPES, ~10 mM $Cl^-$, 4 mM MgATP, pH 7.2, osmolarity adjusted to 345 mmol/kg using sucrose.

pH was adjusted using 1 M NMDG solution. Osmolarity was measured on a vapor pressure osmometer (VAPRO).

#### 2.13.5. Electrophysiology recording

Hi5 cells were seeded onto 10 mm glass coverslips (Ted-Pella, pre-cleaned with 70% ethanol) in 35 mm tissue culture dishes (GenClone 25-200). To express the designs in Hi5 cells, medium was aspirated, and 2 μL of the P2 virus was added to the cells. The dish was put on a rocker at room temperature for 1 hour, and incubated at 27 °C after addition of 2 mL medium. Whole-cell patch-clamp experiments were performed at 18-24 hours after infection. Patch pipettes were pulled from 4 mm thin-wall borosilicate glass capillaries (World Precision Instrument TW150F-4) using Sutter Model P-1000 micropipette puller and had a resistance of 4-7 MΩ when filled with the intracellular solution. Extracellular solutions were applied using a gravity-fed perfusion system equipped with a 8-channel perfusion manifold (Warner Instruments MPP-8), a flow regulating valve (Warner Instruments FR-50), and a cell-bathing chamber (Warner Instruments RC-25). The recording electrode was prepared from silver wires (A-M Systems, 0.008 inch, bare) through electrical chloriding in 3 M KCl solution. A pellet Ag/AgCl electrode (World Precision Instruments, EP2) was used as the reference electrode. Whole-cell currents were recorded using an EPC10 amplifier and PatchMaster Next software (Harvard Biosciences, version 1.2). Current signals were filtered at 2.9 kHz. Cells were patched in the 2 mM [$Ca^{2+}$] extracellular solution. After breaking the patched cell membrane to establish the whole-cell configuration, the bath solution was changed to the 0.02 mM [$Ca^{2+}$] solution. A –100 mV to +100 mV voltage-ramp protocol was applied every 5 seconds with a holding potential of 0 mV. In most cases the recorded currents using this ramp protocol were used to generate the I-V curves. The I-V curve obtained in the 0.02 mM [$Ca^{2+}$] solution was used as the baseline for subtraction of leak current (Extended Data Fig. 6a). The currents measured at –100 mV were used for generating the time courses. After the currents elicited by the voltage-ramp protocol became stable, the bath solution was changed to the one containing the ions of interest (for example, a solution containing 10 mM [$Ca^{2+}$] was used to determine the $Ca^{2+}$ conductance). For measurement of channel conductance for different ions, the 0.02 mM [$Ca^{2+}$] solution was used between the solution exchange to remove the previous ion. Liquid junction potentials between the extracellular and intracellular solutions were calculated to be less than 1 mV (https://swharden.com/LJPcalc/) and therefore was not corrected for the command voltage. The cell capacitance was measured using the automated capacitance correction function of EPC10/Patchmaster Next. Data was exported as ASCII files and analyzed using Python.

#### 2.13.6. Methods for comparing relative conductances of designed channels for different ions

We use Goldman–Hodgkin–Katz (GHK) flux equation to compare the relative conductances of designed channels for different ions based on the measured inward current at –100 mV:

$$\Phi_s = P_s \cdot z_s^2 \cdot \frac{V_m F^2}{RT} \cdot \frac{[S]_i - [S]_o exp(-z_s V_m F/RT)}{1 - exp(-z_s V_m F/RT)} \tag{1}$$

where $\Phi_s$ is the ion flux (i.e., the measured current or current density), $P_s$ is the permeability for ion $S$, $z_s$ is the valence of ion $S$, $V_m$ is the membrane potential, $F$ is the Faraday constant, $R$ is the gas constant, $T$ is the absolute temperature, and $[S]_i$ and $[S]_o$ are the intracellular and extracellular concentrations of ion $S$, respectively.

Given that we are comparing the inward currents and assume that there is no ion of interest inside the cell ($[S]_i = 0$), equation (1) is then simplified to:

$$\Phi_s = -P_s \cdot z_s^2 \cdot \frac{V_m F^2}{RT} \cdot \frac{[S]_o exp(-z_s V_m F/RT)}{1 - exp(-z_s V_m F/RT)} \tag{2}$$

To compare the permeability of two different ions ($S_1$ and $S_2$), Equation (2) can be rewritten as:

$$\frac{P_{s,1}}{P_{s,2}} = \frac{\Phi_{s,1}}{\Phi_{s,2}} \cdot \frac{z_{s,2}^2}{z_{s,1}^2} \cdot \frac{1 - exp(-z_{s,1} V_m F/RT)}{1 - exp(-z_{s,2} V_m F/RT)} \cdot \frac{[S_2]_o exp(-z_{s,2} V_m F/RT)}{[S_1]_o exp(-z_{s,1} V_m F/RT)} \tag{3}$$

Since in most cases the currents at –100 mV are compared ($V_m >> 0$), equation (3) can be simplified to:

$$\frac{P_{s,1}}{P_{s,2}} = \frac{\Phi_{s,1}}{\Phi_{s,2}} \cdot \frac{z_{s,2}^2}{z_{s,1}^2} \cdot \frac{[S_2]_o}{[S_1]_o} \tag{4}$$

To directly compare the relative conductance of the channels for each individual ion using the measured current or current density, the following condition needs to be satisfied:

$$\frac{z_{s,2}^2}{z_{s,1}^2} \cdot \frac{[S_2]_o}{[S_1]_o} = 1 \tag{5}$$

Equation (5) can be written as:

$$\frac{[S_1]_o}{[S_2]_o} = \frac{z_{s,2}^2}{z_{s,1}^2} \tag{6}$$

Therefore, the ion concentration used for determining the relative conductances for different ions was inversely proportional to the square of valence of the particular ion. As a result, the concentration of divalent ions was determined to be 10 mM, and the concentration of $Na^+$ was determined to be 40 mM.

### SI References


56. Sharpe, H. J., Stevens, T. J. & Munro, S. A Comprehensive Comparison of Transmembrane Domains Reveals Organelle-Specific Properties. *Cell* **142**, 158–169 (2010).

57. Levental, I. & Lyman, E. Regulation of membrane protein structure and function by their lipid nano-environment. *Nat. Rev. Mol. Cell Biol.* **24**, 107–122 (2023).

58. Dauparas, J. *et al.* Atomic context-conditioned protein sequence design using LigandMPNN. 2023.12.22.573103 Preprint at https://doi.org/10.1101/2023.12.22.573103 (2023).

59. Jiang, D., Gamal El-Din, T., Zheng, N. & Catterall, W. A. Chapter Five - Expression and purification of the cardiac sodium channel NaV1.5 for cryo-EM structure determination. in *Methods in Enzymology* (eds. Minor, D. L. & Colecraft, H. M.) vol. 653 89–101 (Academic Press, 2021).

60. Jiang, D. *et al.* Open-state structure and pore gating mechanism of the cardiac sodium channel. *Cell* **184**, 5151-5162.e11 (2021).

61. Jiang, D. *et al.* Structure of the Cardiac Sodium Channel. *Cell* **180**, 122-134.e10 (2020).

62. Lenaeus, M., Gamal El-Din, T. M., Tonggu, L., Zheng, N. & Catterall, W. A. Structural basis for inhibition of the cardiac sodium channel by the atypical antiarrhythmic drug ranolazine. *Nat. Cardiovasc. Res.* **2**, 587–594 (2023).

63. Tonggu, L. *et al.* Dual receptor-sites reveal the structural basis for hyperactivation of sodium channels by poison-dart toxin batrachotoxin. *Nat. Commun.* **15**, 2306 (2024).

64. Mastronarde, D. N. SerialEM: A Program for Automated Tilt Series Acquisition on Tecnai Microscopes Using Prediction of Specimen Position. *Microsc. Microanal.* **9**, 1182–1183 (2003).

65. Sun, M. *et al.* Practical considerations for using K3 cameras in CDS mode for high-resolution and high-throughput single particle cryo-EM. *J. Struct. Biol.* **213**, 107745 (2021).

66. Punjani, A., Rubinstein, J. L., Fleet, D. J. & Brubaker, M. A. cryoSPARC: algorithms for rapid unsupervised cryo-EM structure determination. *Nat. Methods* **14**, 290–296 (2017).

67. Sanchez-Garcia, R. *et al.* DeepEMhancer: a deep learning solution for cryo-EM volume post-processing. *Commun. Biol.* **4**, 1–8 (2021).

68. Pettersen, E. F. *et al.* UCSF Chimera—A visualization system for exploratory research and analysis. *J. Comput. Chem.* **25**, 1605–1612 (2004).

69. Croll, T. I. ISOLDE: a physically realistic environment for model building into low-resolution electron-density maps. *Acta Crystallogr. Sect. Struct. Biol.* **74**, 519–530 (2018).

70. Emsley, P., Lohkamp, B., Scott, W. G. & Cowtan, K. Features and development of Coot. *Acta Crystallogr. D Biol. Crystallogr.* **66**, 486–501 (2010).

71. Emsley, P. & Cowtan, K. Coot: model-building tools for molecular graphics. *Acta Crystallogr. D Biol. Crystallogr.* **60**, 2126–2132 (2004).

72. Liebschner, D. *et al.* Macromolecular structure determination using X-rays, neutrons and electrons: recent developments in Phenix. *Acta Crystallogr. Sect. Struct. Biol.* **75**, 861–877 (2019).

73. Williams, C. J. *et al.* MolProbity: More and better reference data for improved all-atom structure validation. *Protein Sci.* **27**, 293–315 (2018).

74. Pettersen, E. F. *et al.* UCSF ChimeraX: Structure visualization for researchers, educators, and developers. *Protein Sci.* **30**, 70–82 (2021).
